# To knot or not to knot: Multiple conformations of the SARS-CoV-2 frameshifting RNA element

**DOI:** 10.1101/2021.03.31.437955

**Authors:** Tamar Schlick, Qiyao Zhu, Abhishek Dey, Swati Jain, Shuting Yan, Alain Laederach

## Abstract

The SARS-CoV-2 frameshifting RNA element (FSE) is an excellent target for therapeutic intervention against Covid-19. This small gene element employs a shifting mechanism to pause and backtrack the ribosome during translation between Open Reading Frames 1a and 1b, which code for viral polyproteins. Any interference with this process has profound effect on viral replication and propagation. Pinpointing the structures adapted by the FSE and associated structural transformations involved in frameshifting has been a challenge. Using our graph-theory-based modeling tools for representing RNA secondary structures, “RAG” (RNA-As-Graphs), and chemical structure probing experiments, we show that the 3-stem H-type pseudoknot (3_6 dual graph), long assumed to be the dominant structure has a viable alternative, an HL-type 3-stem pseudoknot (3_3) for longer constructs. In addition, an unknotted 3-way junction RNA (3_5) emerges as a minor conformation. These three conformations share Stems 1 and 3, while the different Stem 2 may be involved in a conformational switch and possibly associations with the ribosome during translation. For full-length genomes, a stem-loop motif (2_2) may compete with these forms. These structural and mechanistic insights advance our understanding of the SARS-CoV-2 frameshifting process and concomitant virus life cycle, and point to three avenues of therapeutic intervention.

## Introduction

While the novel coronavirus agent, SARS-CoV-2, has decimated world economies, influenced political leadership, infected more than 180 million people, and claimed the lives of 3.9 million, the level of scientific cooperation and advances we have witnessed this past year is remarkable. Besides successful vaccine development efforts, progress on unraveling the complex and multifarious biophysical aspects of the virus life cycle and infection trajectory has helped us describe how the virus hijacks our own protein-synthesis machinery into making viral proteins efficiently and propose new lines of defense against the deadly disease it carries. These insights about the life cycle of the virus and mode of action are invaluable for further development of drugs and other strategies to combat future viral epidemics.

Although viral proteins have been a focus of many scientific groups, investigations of the RNA viral agent itself are crucial for understanding how the RNA invader replicates itself, is translated by the human ribosomal machinery, assembles, and synthesizes a suite of viral proteins that enable the continuation of its invasive trajectory. RNA-targeting therapeutics and vaccines can disarm the origin of the infection rather than its products and be more effective in the long term. However, the complexity of the RNA molecule and the lagging science about its modeling, imaging, and drug screening compared to proteins pose challenges. With technological improvements in RNA delivery systems, the rise of CRISPR-based gene editing systems, ^1^ and improved RNA modeling techniques, ^2,3^ this RNA focus is not only warranted but clearly successful, as evident by recent vaccines.

Of particular interest by many groups is the RNA frameshifting element (FSE), a small region in the open reading frame ORF1a,b region (Fig. 1, top) of the viral genome that codes for the polyproteins that initiate the cascade of viral protein synthesis. The FSE is responsible for the crucial −1 programmed ribosomal frameshifting (−1 PRF) mechanism utilized by many viruses including HIV-1 to handle protein synthesis from overlapping reading frames. ^4–6^ Its stimulatory pseudo-knot or stem-loop motif is believed to be crucial for the requisite pausing. ^6–10^ When encountering ORF1b, out of register with respect to ORF1a, the ribosome backs up one nucleotide in the 5′ direction to define a different sequence of codons (Fig. 1). Given noted correlations between the conformational plasticity of the stimulatory element and frameshifting efficiency, more complex pausing mechanisms may be involved than a simple “barrier”. ^11–14^

**Figure 1:**
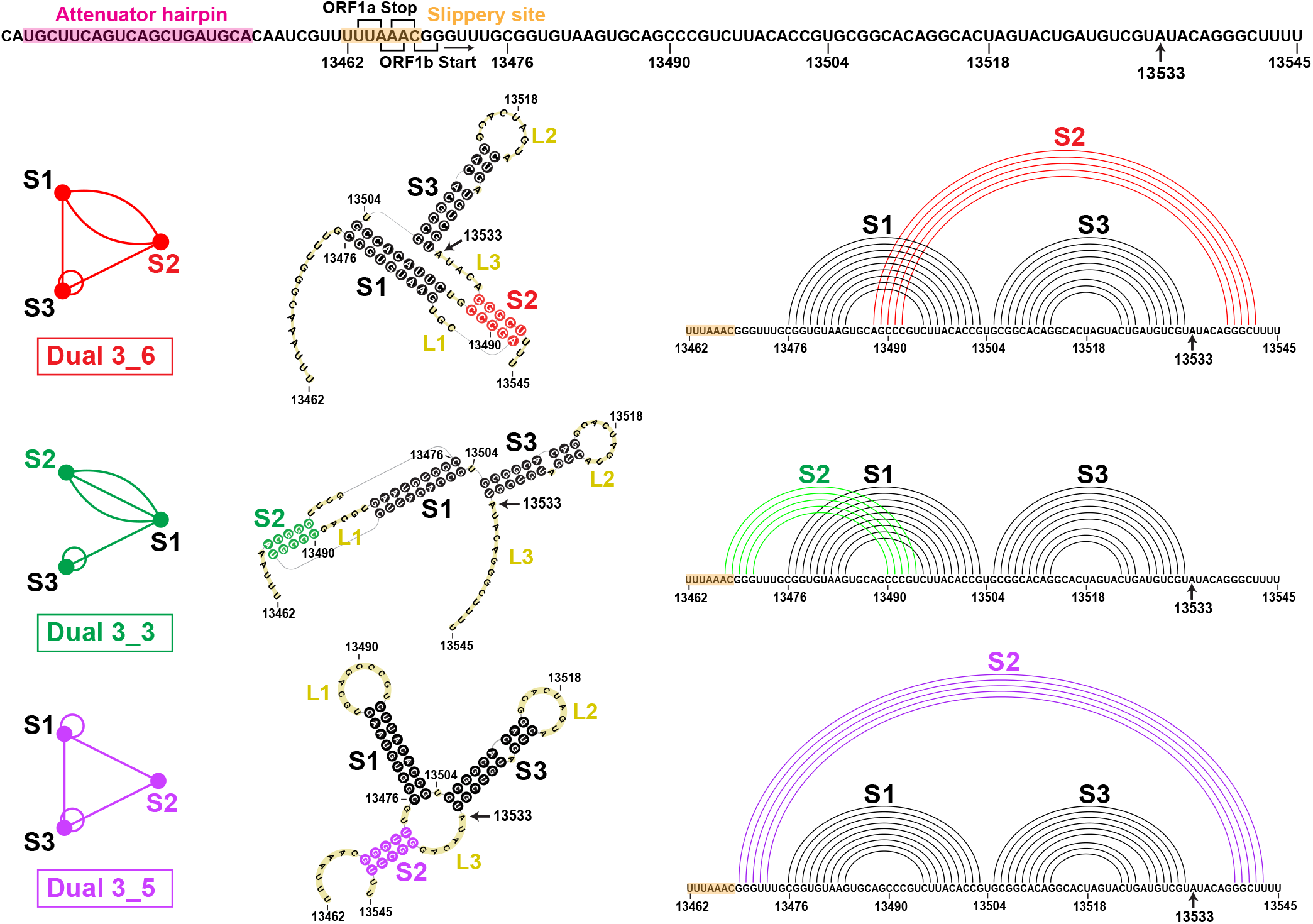
The FSE sequence and three relevant 2D structures for the SARS-CoV-2 84 nt frameshifting element (residues 13462–13545) emerging from this work that combines 2D structure prediction, SHAPE structural probing, and thermodynamic ensemble modeling. The −1 frameshifting alters the transcript UUU-UU*A(Leu)-AAC(Asn)-GGG at the second codon (asterisk) to backtrack by one nucleotide and start as AAA-CGG(Arg) instead, so that translation resumes at CGG. At top is the FSE sequence, with the attenuator hairpin region and the 7 nt slippery site highlighted and A13533 labeled (C in SARS). The ORF1a end and ORF1b start codons for the overlapping regions are marked. For each 2D structure, H-type 3_6 pseudoknot, HL-type 3_3 pseudoknot, and three-way junction 3_5 (unknotted RNA), corresponding dual graphs, 2D structures, and corresponding arc plots are shown, with color coded stems and loops labeled.

The 84-residue SARS-CoV-2 FSE (13462–13545 of the 29891 nt RNA genome) contains a 7-residue slippery site (UUUAAAC) and a 77-residue RNA (Fig. 1). An upstream attenuator hairpin (Fig. 1) may play a role in frameshifting. ^15,16^ The FSE’s crucial role in viral protein synthesis makes it an excellent target for therapeutic intervention. ^14,17,18^ Indeed, small-molecule agents such as *1,4-diazepane derivative 10* (MTDB) (originally designed for SARS-CoV ^13,15,19^), fluoroquinolone antibacterial *merafloxacin*, ^20^ and a phenyl thiourea C5^16^ were found to hamper SARS-CoV-2 frameshifting.

Because of the crucial relationship between the FSE conformational plasticity and the frameshifting mechanism, it is important to unravel the FSE conformational landscape. Complex interactions are likely involved both within the FSE and between the FSE and the ribosome. Here we focus on better understanding of this FSE conformational landscape using a combination of complementary graph-based modeling and chemical reactivity experiments. Already, several groups have explored FSE structure by modeling, ^21–25^ *in vivo* Selective 2′-Hydroxyl Acylation by Primer Extension (SHAPE) ^26,27^ and DMS structure probing experiments, ^20,28–34^ NMR, ^35^ Cryo-EM, ^29,36^ and other biophysical mutational profiling and scanning experiments. ^15,37,38^ Many have characterized the FSE as a 3-stem H-type pseudoknot with colinear Stems 1 and 2 intertwined via a pseudoknot and perpendicular Stem 3. This association has persisted from the SARS-CoV FSE characterization, ^15^ which differs in only one base from the SARS-CoV-2 FSE (residue A13533 in Covid-19 is C in SARS, Fig. 1). However, depending on the modeling software and experimental technique, alternative secondary structures have been offered for SARS-CoV-2, both pseudoknotted and unknotted (see below). ^20,23–25,28–34^

In our prior work ^22^ (see also commentary^39^), we defined target residues for drug binding and gene editing of the FSE from designed minimal mutants that dramatically transform the FSE conformation. Our RAG (RNA-As-Graphs) machinery represents RNA 2D structure as coarse-grained dual graphs, where double-stranded RNA helices are represented as vertices, and loop strands are edges. The advantage of graphs is that they are robust and capture the topology of the RNA while allowing for differences in the lengths of stems and loops; thus, the same graph corresponds to multiple 2D models that differ in sizes of stems and loops. This makes structure comparison, transformation, and design more facile and efficient. The common H-type pseudoknotted structure of the FSE corresponds to the 3_6 dual graph in Fig. 1. Using our RAG-based genetic algorithms for RNA design by inverse folding, ^40^ we designed mere double mutants that transformed the 3_6 conformation for a 77 nt FSE (no slippery site) (Fig. 1) into 3-stem and 2-stem structures with and without pseudoknots. Microsecond molecular dynamics simulations of these mutants modeled at atomistic detail with explicit solvent demonstrated the stability of these alternative forms. Among these mutants, the 3-way junction (dual graph 3_5) is further investigated here.

We also highlighted how structure predictions of the FSE by available programs depend on the length considered. ^22^ The lengths are both computationally and biologically meaningful, since the slippery site is thought to be inaccessible while the FSE is in direct interaction with the ribosome, but possibly free otherwise. Besides the slippery site, neighboring units, especially the up-stream nucleotides, also influence the predicted topologies. We showed that the sequence context of 77, 84, and 144 nt leads to various structure predictions for the FSE that are both pseudoknotted and unknotted (Fig. 3 here too). ^22^

Here, we continue to untangle this length dependence through graph theory modeling combined with SHAPE experiments. Our combined analysis describes a conformational landscape with *three viable structures* of the FSE: two pseudoknotted RNAs (3_6 and 3_3 in our dual graph notation, or H-type and HL-type 3-stem pseudoknots), and one unknotted, 3-way junction RNA (3_5) (Fig. 1).^*^ The flexible Stem 2 may be involved in a switch between these conformations and associations with the ribosome during protein translation, as well as define a co-transcriptional kinetic folding trap. For whole genome constructs, a stem-loop motif may compete with these forms. Thus, our mutants which stabilize one form over the others may be particularly effective when used in combination with anti-viral therapy that targets a specific FSE form.

We first examine sequence and structure conservation of the FSE region in coronaviruses and current SARS-CoV-2 variants, and highlight length-dependent predictions of 2D FSE structures. Second, we present results guided by SHAPE reactivities that point to two pseudoknots and one 3-way junction 2D topologies. Third, we predict and experimentally confirm mutants that strengthen each of these three conformations, and present a predicted conformational landscape for FSE RNAs of length 77 to 144 nt. Fourth, we discuss other 2D FSE structures in the literature, probe alternative forms for longer genome contexts, compare reactivity data to date, and follow with some computational mutations motivated by Bhatt et al. ^36^

Together, the SHAPE data and statistical landscape modeling help describe the relation between FSE length and structure, as well as implications to frameshifting mechanisms involving the ribosome. These results help consolidate FSE reports to date and define new therapeutic avenues for regulating frameshifting efficiency and hence Covid-19 infections.

## Results

### Multiple sequence alignment and variant analysis of coronaviruses emphasize FSE features

To put into context the FSE structure of SARS-CoV-2 and pinpoint the relative flexibility of the different stems, we analyze the sequence similarity of an enlarged FSE region of 222 nt (residues 13354–13575) in the coronavirus family by multiple sequence alignment (MSA). Among 1248 non-redundant coronavirus sequences downloaded from Virus Pathogen Database and Analysis Resource (ViPR), ^41^ 182 non-duplicate homologous sites are structurally aligned to the SARS-CoV-2 FSE using the Infernal covariance model ^42^ (see Methods). We show the alignment for 16 top scored coronaviruses in Fig. 2. For each virus, genome identity with the entire SARS-CoV-2 and for only the 222 nt FSE are indicated. Darker purple shadings indicate greater sequence homology.

**Figure 2:**
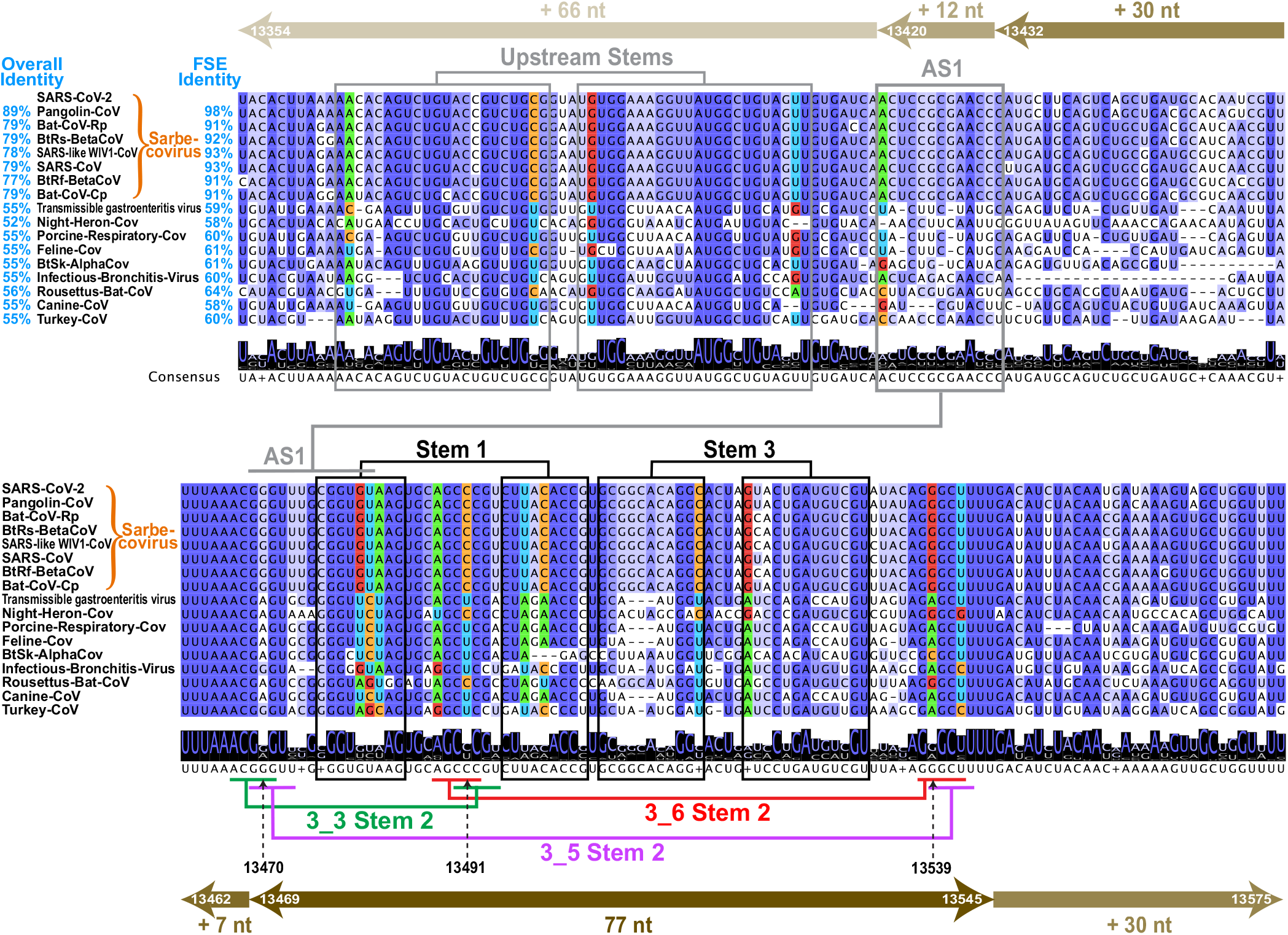
Multiple sequence alignments (MSA) of coronavirus frameshifting elements found by the Infernal covariance model ^42^ shown for 16 top-scored sequences among 182 unique homologues. Arrows at top and bottom illustrate the FSE expansion from 77 to 84 nt (+7 slippery site nt), 144 nt (+30 nt on both ends), 156 nt (+12 upstream nt), and 222 nt (+66 upstream nt). Sixteen top scored coronaviruses are aligned with the SARS-CoV-2 222 nt FSE region (insertions are hidden), with sequence similarities shown for both the whole genome and the FSE region. Nucleotides are colored based on sequence conservation. The consensus sequence is written below with a sequence logo (at each position, the overall stack height indicates sequence conservation level, and the height of an individual letter within indicates the relative frequency of that nucleotide). Stems are marked based on our analysis here: black for Stems 1 and 3, red/green/purple for Stem 2 of 3_6/3_3/3_5, consistent with Fig. 1, and grey for Alternative Stem 1 (AS1) and upstream stems. The covarying base pairs detected by R-scape ^43^ are colored by nucleotide identity: green A, blue U, orange C, and red G.

In the central 84 nt FSE region (residues 13462– 13545) corresponding to Fig. 1, consensus Stems 1 and 3 are colored black, with Stem 2 corresponding to the H-type pseudoknot (dual graph 3_6) in red, HL-type pseudoknot (3_3) in green, and 3-way junction (3_5) in purple. We see that Stem 1 is highly conserved, with deletions in the 3′ strand in only one distant coronavirus. Moreover, subsequent covariation analysis using R-scape ^43^ (see Methods) detects 2 strong covarying base pairs (colored by nucleotide in Fig. 2, i.e., green A, blue U, orange C, and red G), also found in the phylogenetic analysis by Andrews et al. ^24^ Many deletions are found in Stem 3 and the sequences are less conserved, suggesting different locations and lengths for Stem 3 in different coronaviruses. Stem 2 is the shortest and the most flexible. In SARS-CoV-2, the two pseudoknots 3_6 and 3_3 have equally strong Stems 2, both made of one AU and four GC base pairs. The two Stems 2 share the same central CCC region (13490–13492) but involve different base pairing orientations (Stem 1 loop base pairs with the 3′ end in 3_6, and with the 5′ end in 3_3). While these two Stems 2 are fully conserved in Sarbecovirus subgenus, the middle C13491 in the shared region is mutated to U in more distant coronaviruses. Interestingly, some compensatory mutations from G to A occur at complementary locations for both 3_6 (residue 13539) and 3_3 (13470) Stem 2. While this compensatory mutation in 3_6 Stem 2 is considered a covariation by R-scape, some G13470A mutations in 3_3 Stem 2 occur without the C13491U mutation, resulting in the A13470-C13491 mismatch in some non-Sarbecovirus sequences, which suggests that the 3_3 Stem 2 is Sarbecovirus-specific. Stem 2 of the 3_5 junction is less stable, made of one GC and four GU base pairs.

By extending the upstream sequence, a stem-loop Alternative Stem 1 (AS1) competing with Stem 1, and Stem 2 of 3_3 and 3_5 emerges. This AS1 appears in several groups’ whole genome chemical probing, ^20,30–33^ and is also predicted by our SHAPE probing here for 156 and 222 nt constructs (see Comparison section). We see that AS1 is only conserved in Sarbecoviruses, with many deletions in the 5′ strand in distant coronaviruses, and only a weak covarying base pair is found. Therefore, both sequence conservation and covariation analysis suggest that Stem 1 is most conserved in the coronavirus family, while 3_3 Stem 2 and AS1 may be Sarbecovirus-specific.

To demonstrate the sequence conservation of the SARS-CoV-2 FSE RNA, we analyze 459421 variants of SARS-CoV-2 deposited on GISAID (Global Initiative on Sharing All Influenza Data) database ^44^ by February 12, 2021 (Fig. S1). Only 8504 or 2% exhibit mutations in the FSE segment. Among the mutated sequences, 98% are single mutants. Mutation maps for the 84 nt FSE and the spike gene segment per nucleotide (plotted on different scales) show that the spike gene region has an order of magnitude more mutations than the FSE. Interestingly, residue A13533, which is C in SARS-CoV, is never mutated to C, but only to G. Further analysis of recent highly transmissible British (B.1.1.7), South Africa (B.1.351), Brazil (P.1), and New York City (B.1.526) Covid-19 variants also show concentrated mutations in the spike gene region, with 4-12 residues having mutation rates > 85%, and very few (random) mutations in the FSE (Fig. S2). This analysis reinforces the high conservation of the FSE region and its suitability for anti-viral therapy, consistent with other sequence variation studies. ^45^

### Length dependent RNA 2D structure predictions raise caveats

Several works have scanned experimentally RNA genomes with windows of variable lengths. ^24,37,46^ In secondary structure predictions of RNAs, 120 nt is considered reasonable for predictions. ^47,48^ Indeed, in our application of five 2D folding programs that can predict pseudoknots (PKNOTS, ^49^ NUPACK, ^50^ IPknot, ^51^ ProbKnot, ^52^ and vsfold5^53^) to four RNAs with pseudoknots, we find that the 120 nt window recommended in the literature appears reasonable in general (Fig. S3). For the FSE, we extend our length-dependent predictions ^22^ using PKNOTS, NUPACK, IPknot, and ProbKnot to generate optimal 2D structures for 4 lengths: 77, 84, 144, and 156 nt (see Fig. 2). Fig. 3 shows the 2D arc plots for hydrogen bonding, along with the associated dual graphs for corresponding optimal structures.

**Figure 3:**
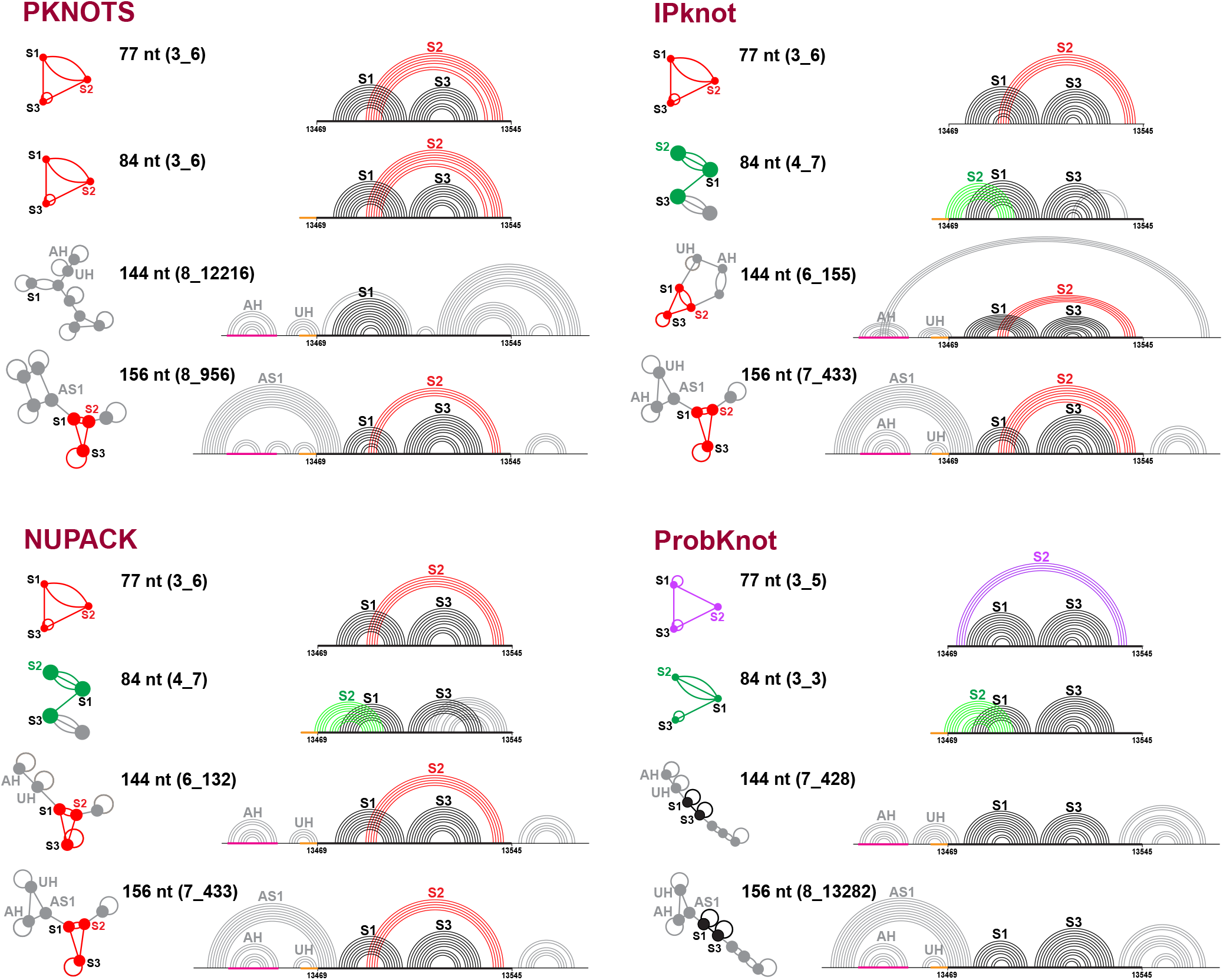
Predicted optimal structures for the frameshifting element using PKNOTS, NUPACK, IPknot, and ProbKnot (see text). For each program, 4 different sequence lengths are used: 77, 84, 144, and 156 nt. The common 77 nt subsequence is aligned, the slippery site is colored orange, and the attenuator hairpin AH is magenta. The predicted structures are shown as arc plots, with Stems 1 and 3 in black, and Stem 2 of 3_6, 3_3, and 3_5 in red, green, and purple, respectively. An upstream hairpin that blocks 3_3 Stem 2 and Alternative Stem 1^28–31^ are labeled UH and AS1, respectively. Corresponding dual graphs 3_6 (red), 3_3 (green), 3_5 (purple), and 2_1 (black) are highlighted as graphs or subgraphs of larger motifs.

We see that for 77 nt, 3 out of the 4 programs predict a 3_6 pseudoknot (H-type), but ProbKnot predicts the 3_5 3-way junction. For 84 nt, only PKNOTS predicts the 3_6 pseudoknot, while ProbKnot predicts the 3_3 pseudoknot (HL-type), and IPknot and NUPACK predict a two-pseudoknot structure 4_7. This 4_7 graph can be partitioned, using our partition algorithm for dual graphs, ^54^ into subgraphs 3_3 and 2_3 (see Fig. S4), with the former corresponding to the 3_3 pseudoknot, and the latter to the new pseudoknot formed by the 3′ end intertwining with Stem 3. Stem 2 of the 3_3 pseudoknot contains 7-9 base pairs and involves 2 residues in the slippery site, which explains why it does not appear in the 77 nt system.

For 144 nt FSE, the predictions are quite different. Only Stem 1, the attenuator hairpin AH, and an upstream hairpin UH which blocks the 3_3 Stem 2 are consistently predicted. Both IPknot and NUPACK predict a 3_6 pseudoknot in the central 77 nt FSE region, but IPknot binds the 3′ end with the 5′ end hairpin loop to form another pseudoknot (6_155), while NUPACK predicts a 3′ end hairpin (6_132). ProbKnot only predicts Stems 1 and 3 in the central 77 nt, which corresponds to a 2_1 dual graph.

For 156 nt, the 3_6 pseudoknot recurs (only ProbKnot predicts a 2_1), and the Alternative Stem 1 (AS1) appears in all four predictions. However, both AS1 and Stem 1 co-exist in our systems. Others found that an extended AS1 can exclude Stem 1 and result in a unknotted structure with only 3_6 Stems 2 and 3 (2_2). ^29–31^ The AS1 together with the attenuator hairpin and stem UH can form an upstream 3-way junction, which blocks Stem 2 of 3_3 and 3_5.

These 2D predictions show a strong dependence of the FSE structure on sequence length, and underscore how the 77 nt central region can form alternative stems with upstream sequences. Both multiple sequence alignment and length-dependent predictions show that Stem 1 is highly conserved, while Stem 2 is variable for this length (Fig. 2).

### SHAPE reactivity data reveals dominant alternative pseudoknot in longer sequence contexts and minor 3-way junction

To experimentally probe the formation of alternative structures in the SARS-CoV-2 FSE, we investigate the SHAPE reactivity of two RNA FSE constructs of 77 nt (residues 13469–13545) and 144 nt (residues 13432–13575) in Fig. 4.

**Figure 4:**
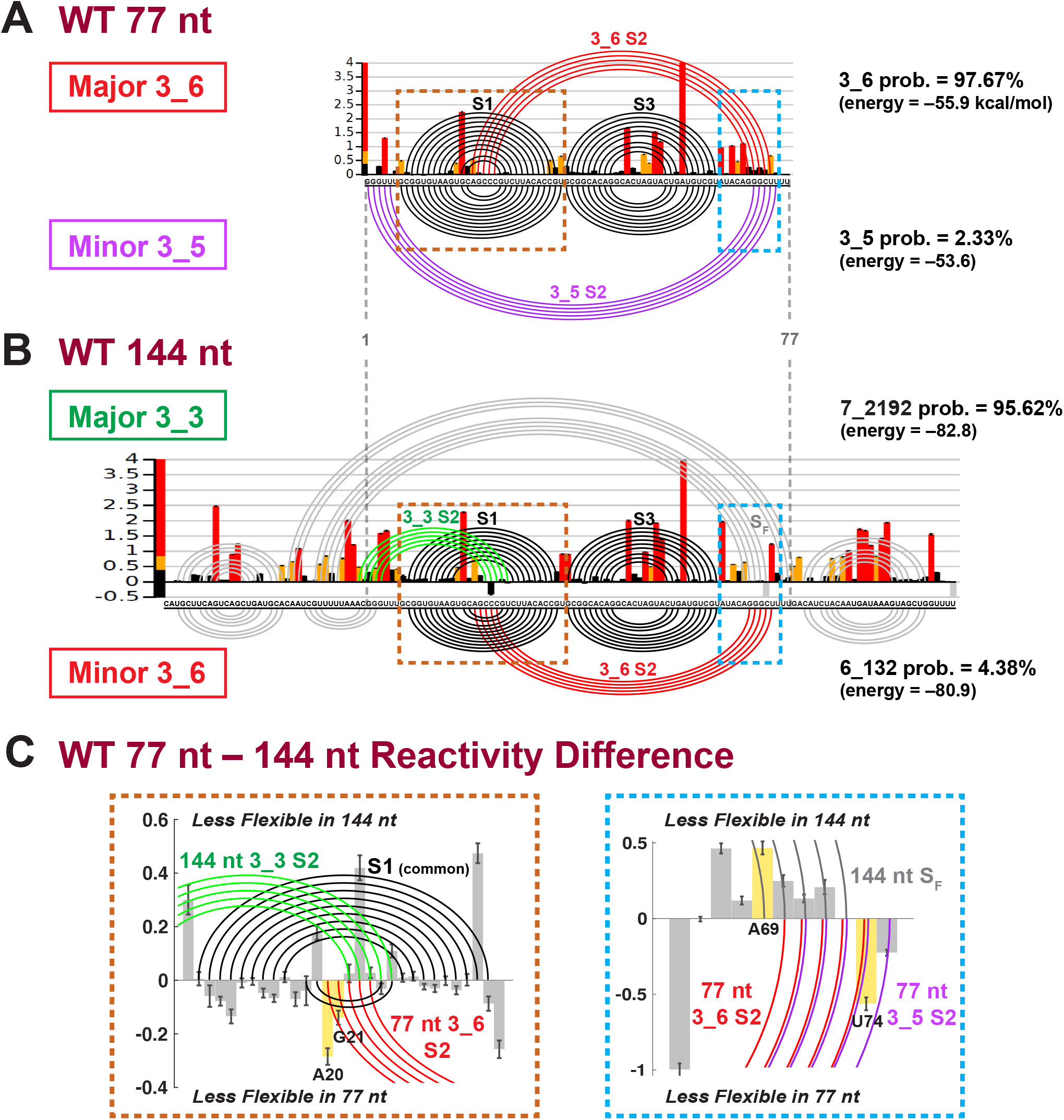
SHAPE reactivity analysis for SARS-CoV-2 frameshifting element for 77 and 144 nt (Replicate 1). (A) The SHAPE reactivity for the 77 nt construct is plotted by bars, with red/yellow/black representing high/medium/low reactivity. The arc plot at top shows the dominant 3_6 pseudoknot predicted by the ShapeKnots energy landscape (98% of conformational space), and at bottom is the minor 3_5 (2% of landscape). Stems are labeled, and the Gibbs free energy (*kcal/mol*) and Boltzmann distribution probabilities are given. (B) SHAPE reactivity and ShapeKnots predictions for 144 nt construct. (C) Reactivity differences between the two constructs are shown for two enlarged key regions highlighted in A,B, with positive/negative differences indicating less flexibility in the 144/77 nt construct. Base pairs in the 144 nt 3_3 conformation are plotted by arcs at top, and 77 nt 3_6 or 3_5 at bottom. Critical residues for reactivity comparisons are highlighted.

In general, SHAPE experiments provide structural anchors for interpreting RNA structures by exploiting the high reactivity of free 2′-hydroxyl groups of the RNA ribose sugar to suitable chemical reagents. The measured reactivities at each nucleotide are directly correlated to the local RNA flexibility, and the paired bases will generally have low reactivity. These experimental data are used to define modified base pair probabilities that guide the energy minimization in the structure prediction program (ShapeKnots). ^55,56^

For each FSE length, we probed two replicates, with 5NIA reagent and Bicine buffer (see Methods, SI for alignments of replicates, and Table 1). In Fig. 4, the SHAPE reactivities of Replicate 1 are shown as histograms plotted per residue, and arc plots above correspond to the dominant prediction. Arc plots below the reactivity data correspond to minor conformers. Because ShapeKnots predicts multiple structures ranked by free energies (not just a minimum free-energy structure), we apply Boltzmann weighting (*p_i_* = exp (−*E_i_* / (*k_B_* T)) where *p_i_* and *E_i_* are the probability and free energy for conformer *i*, *k_B_* is the Boltzmann constant, and T is room temperature, set at 37 °C) to calculate the energy landscape contribution of each conformer.

**Table 1:**
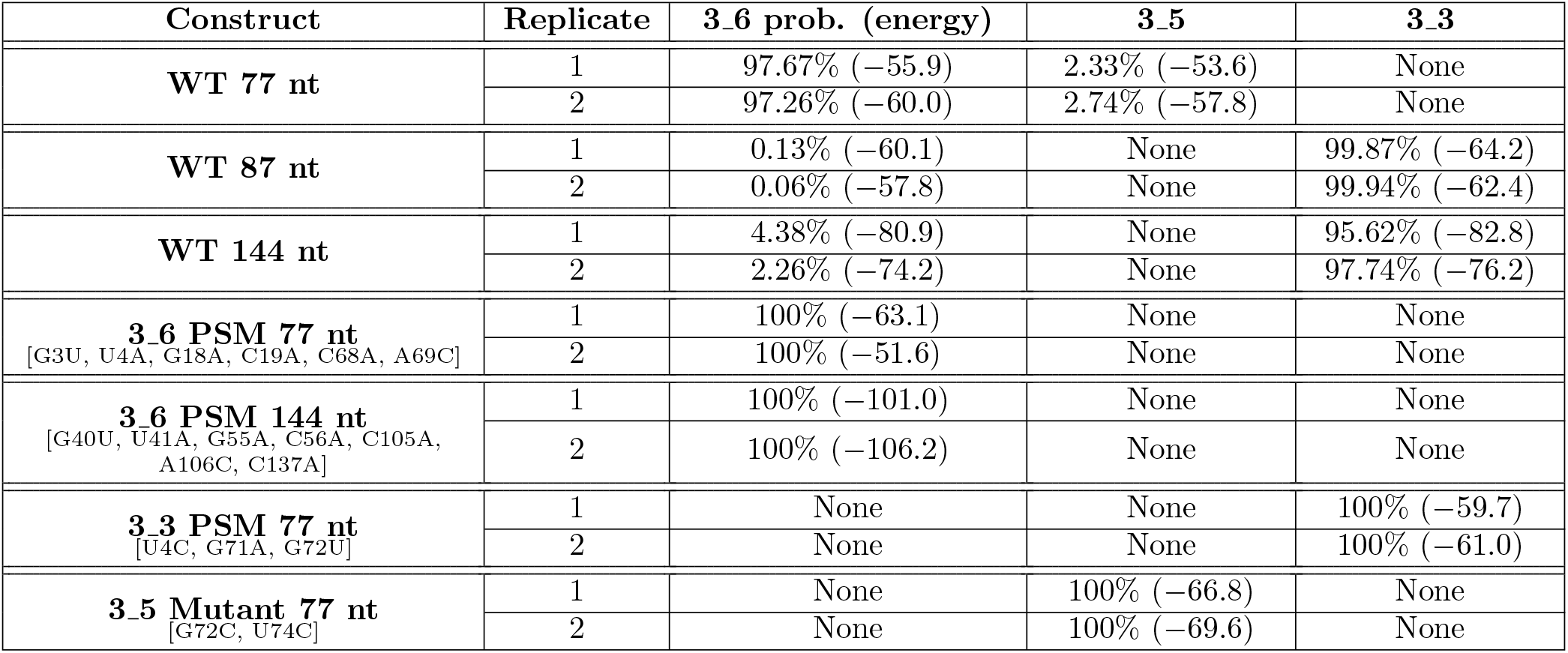
Summary of ShapeKnots prediction results for wildtype frameshifting element and mutants developed and tested in this work. For each construct, the probability and free energy (*kcal/mol*) predicted for 3_6 pseudoknot, 3_5 junction and 3_3 pseudoknot are shown. The mutations are annotated by their positions in the relative constructs; 77 nt construct covers residues 13469–13545, 87 nt covers 13459–13545, 144 nt covers 13432–13575. PSM: Pseudoknot-strengthening mutant; see next section.

Consistent with the modeling for 77 nt, when we incorporate its SHAPE reactivity in ShapeKnots 2D structure ensemble predictions, we find that 98% of structures form the 3_6 pseudoknot, with the 3_5 3-way junction playing a minor role (Fig. 4A). The 3_6 pseudoknot has the same structure as the predictions by PKNOTS, IPknot, and NUPACK (Fig. 3), with small variations in stem lengths. The 3_5 junction nevertheless has a shifted Stem 2 towards the 5′ and 3′ end, compared with the prediction by ProbKnot.

The same experiment on the 144 nt construct detects the 3_6 pseudoknot only in 4.4% of the population, while the 3_3 pseudoknot represents 95.6% of the landscape (Fig. 4B). The 144 nt 3_6 conformation agrees with NUPACK’s prediction in Fig. 3 (dual graph 6_132). Comparing the two pseudoknotted structures (top versus bottom arc plots), we see that Stems 1 and 3 are very similar, but Stem 2 is different. In 3_3, the pseudoknot involves Stem 1 intertwining with Stem 2 at the 5′ end of the FSE, while for 3_6, Stem 1 loop region hydrogen bonds with the 3′ end of the FSE to form Stem 2 (Fig. 1).

The computed difference in SHAPE reactivity (77 nt reactivity minus 144 nt reactivity) reveals changes consistent with these findings in two key regions (Fig. 4C): Stem 1 with its loop and the 77 nt 3′ end. In the Stem 1 loop, two residues A20 and G21 (numbered in the 77 nt context, equivalent to A13488 and G13489) are only paired in the 3_6 Stem 2, and are more flexible in the 144 nt system. Commensurately, the complementary residue U74 of A20 on the 3′ end also has increased flexibility in 144 nt. Similarly, for 144 nt, Stem *S_F_* flanking the 3_3 pseudoknot involves a critical residue A69 (less flexible in 144 nt) absent from base pairs associated with the 3_6 pseudoknot and 3_5 junction.

Replicate 2 (Fig. S5) reaffirms our finding of a dominant 3_6 pseudoknot (97%) and a minor 3_5 junction for 77 nt. For 144 nt, besides the 3_3 pseudoknot conformation (dual graph 7_2192) that was dominant in Replicate 1, Replicate 2 yields another 3_3-containing structure (dual 6_383). The two structures share the central 3_3 pseudoknot of Replicate 1 but differ in the flanking regions. Namely, in 57% of the conformations, *S_F_* is replaced by a hairpin at the 3′ end (see Fig. S5). In Fig. S6A, we see that the two aligned replicates for each construct agree well with one another, especially for the 77 nt construct.

We also used dimethyl sulfate (DMS) chemical probing coupled with mutational profiling to identify correlations in the structure. The PairMap technique identifies correlation ^57^ to suggest not only which residues are paired but with whom they may pair. In Fig. S6B, the dark arcs from PairMap indicate the principal interactions, while the lighter colored arcs correspond to minor interactions. Consistent with the multiple sequence alignment (Fig. 2), Stems 1 and 3 (for 77 nt) and Stem 1 (for 144 nt) are strongly preserved, while Stem 2 is more tentative. The minor Stem 2 for both lengths corresponds precisely to the Stem 2 in the two pseudoknotted structures above. This additional experimental approach supports our findings and is based on a direct analysis of DMS mutational reactivities, independent of thermodynamic modeling.

### Mutant predictions for dominating the conformational landscape by 3_6, 3_3, and 3_5 topologies are confirmed by SHAPE

Our SHAPE and correlated DMS experiments suggest three relevant structures that make up the FSE conformational landscape for lengths up to 144 nt: two pseudoknots (3_6 and 3_3), and a 3-way junction (3_5). As this conformational flexibility may play a mechanistic role in frameshifting, we sought to stabilize each conformer by minimal mutations. Such analysis can aid anti-viral therapy by suggesting how to target a specific FSE conformer and also provides insights into possible transitions between the three conformers.

We apply our RAG-based software RAG-IF ^40^ as developed and applied in our prior work ^22^ to determine minimal mutations for each conformer to dominate the landscape. Briefly, our genetic algorithm works by transforming one dual graph into another by iterating on a sequence of mutations in pre-selected regions so as to minimize the difference (measured by Hamming distance) between the current and target graph, in the spirit of a natural selection process; the fold of each graph is determined by a consensus between two 2D folding programs. See Methods and Refs. ^40^

To design the 3_6 pseudoknot-strengthening mutant (PSM), we apply RAG-IF to transform the 3_5 predicted by ProbKnot (Fig. 3) onto 3_6 (Fig. 5A). A 4-residue mutant [G3U, U4A, C68A, A69C], which breaks Stem 2 of 3_5 and creates two extra base pairs for Stem of 3_6, is selected for the 77 nt construct. With additional 2D prediction program screening (see Fig. S7A), we add two mutations [G18A, C19A] to further strengthen Stem 2 with 4 additional base pairs. For 144 nt, after testing the above 6 mutations using four 2D prediction programs (Fig. S7B), we add a mutation to the 3′ end to inhibit a stem that interferes with Stem 2 of 3_6. The resulting 7-residue mutant is [G40U, U41A, G55A, C56A, C105A, A106C, C137A].

**Figure 5:**
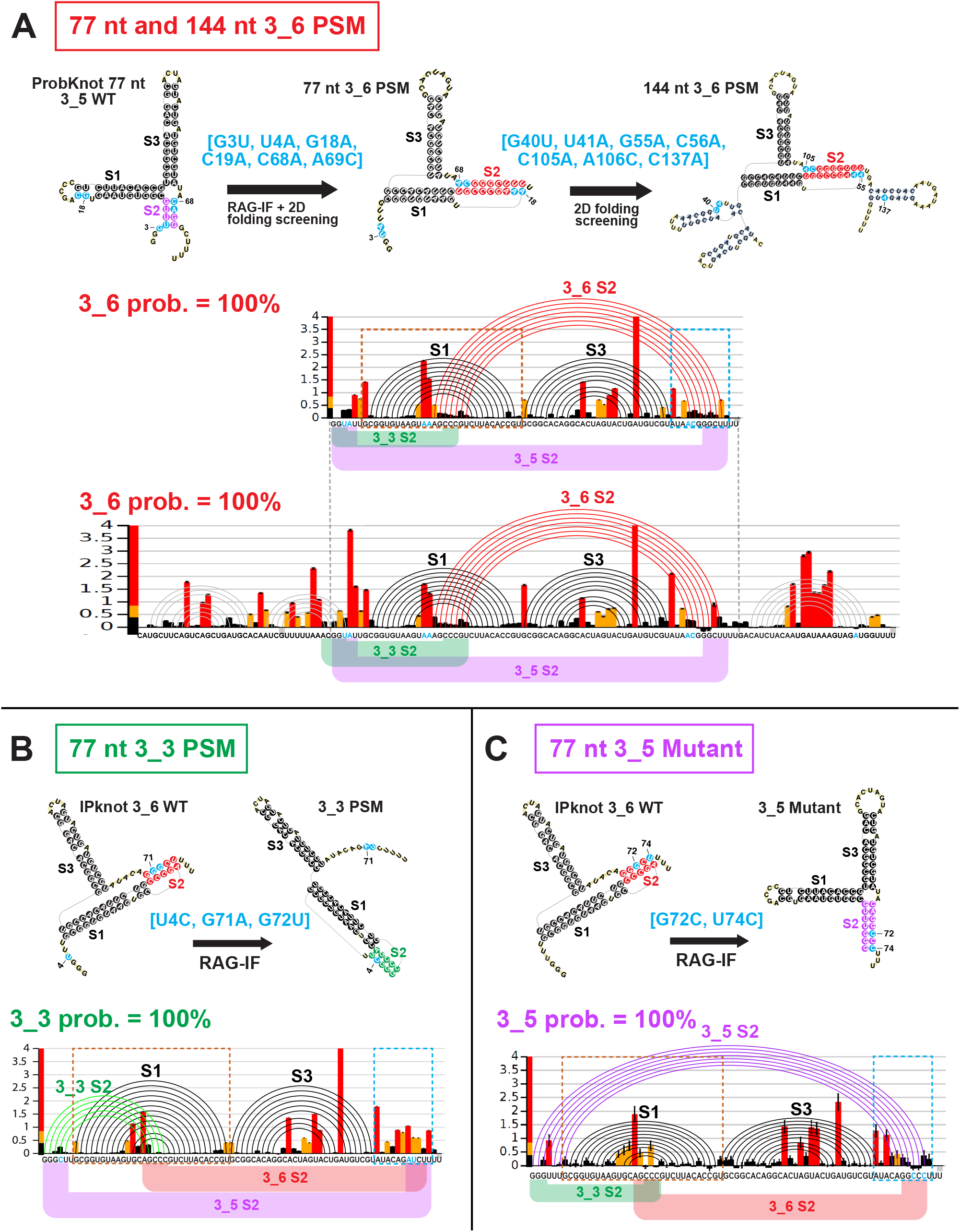
Design and SHAPE analysis for (A) 77 and 144 nt 3_6 pseudoknot-strengthening mutant (PSM), (B) 77 nt 3_3 PSM, and (C) 77 nt 3_5 Mutant. For each mutant, we show the design flow, where we use RAG-IF and multiple 2D structure prediction program screening to determine mutations that stabilize the 3_6 pseudoknot, 3_3 pseudoknot, or 3_5 junction. Mutations are highlighted in blue. The SHAPE reactivity bar plots, and arc plots of the structure predicted by ShapeKnots, with alternative Stem 2 positions are shown. See Fig. S8 for reactivity differences between the mutants and the wildtype for two boxed key regions.

Subsequent SHAPE experiments confirm our predictions for both the 77 and 144 nt constructs of this 3_6 PSM: 100% of the landscape is now occupied by the 3_6 pseudoknot (Fig. 5A), when chemical reactivity data are incorporated in the 2D structure prediction by Shape-Knots. The 3_6 Stem 2 has 7 instead of the expected 9 base pairs, but is longer than the wildtype Stem 2 (Fig. 4A). Moreover, when we compare the 3_6 PSM with the wildtype for 77 nt constructs (Fig. S8A), the reactivity differences support the two new base pairs in Stem 2. Residue G25 was base paired with C19 in the wildtype FSE, but after the C19A and A69C mutations, it pairs with 69C. As a result, C19 has increased flexibility and 69C shows decreased flexibility in the PSM SHAPE data. Similarly, for the new base pair U26 with 68A, we note decreased reactivity for 68A.

We also design a 3_3 pseudoknot-strengthening mutant for the 77 nt FSE similarly. We choose a triple mutant [U4C, G71A, G72U] that is predicted to form 5-7 base pairs for 3_3 Stem 2 (see Fig. S9 and Methods). Subsequent SHAPE reactivities in Fig. 5B show that the conformational landscape is now 100% 3_3 pseudoknot. The comparison between the 3_3 PSM and the wildtype 77 nt reactivities (Fig. S8B) show very small differences in the Stem 1 region, because Stem 2 of the wildtype 3_6 and the PSM 3_3 overlap. Nevertheless, we see increase in flexibility at the 3′ end, where the 3′ strand of Stem 2 in 3_6 and 3_5 locate, supporting 3_3 pseudoknot over the other two conformations in this predicted mutant.

Because the unknotted 3-way junction (3_5) emerges as a minor player in the 77 nt FSE conformational landscape, we obtain reactivity data for the double mutant we had predicted in our prior work ^22^ to stabilize this fold over the two pseudoknots. Fig. 5C shows that merely two mutations [G72C, U74C] on the 3′ edge of 3_6 Stem 2 accomplish this dramatic change. This 3-way junction becomes the sole conformer in the 77 nt mutant landscape, compared to 2–3% in the wildtype (Fig. 4A). By examining reactivity differences with the wildtype 77 nt (Fig. S8C), we find that residues in the loop region of Stem 1, which are base paired in the wild-type 3_6 Stem 2, become more flexible in this mutant. Moreover, a 3′ end residue A69, which is newly base paired in this mutant’s Stem 2, has decreased reactivity, again supporting the 3_5 conformation.

The combined evidence points to a conformational landscape for the SARS-CoV-2 FSE that is sensitive to the sequence length and highlights two major players — an H-type pseudoknot (3_6 dual graph) and an HL-type pseudoknot (3_3 dual graph) — as well as a minor 3-way junction (unknotted) (3_5). Our mutant predictions for strengthening all three structures are confirmed by two SHAPE replicates (see Table 1 and Fig. S10 for alignments). Although the 3_6 pseudoknot (98%) dominates the wildtype 77 nt landscape, with only 2 or 3 mutations, we can shift the landscape to be 100% 3_5 3-way junction or 100% 3_3 pseudoknot. Hence, all 3 conformations are viable for the FSE, and they may not be too far away from one another from a sequence landscape point of view.

### Consensus conformational landscape of the FSE clarifies length dependence

To consolidate the above information, we estimate the energy landscape for the three viable conformations as a function of sequence length. We consider two ways of expanding the FSE sequence: (1) asymmetrically adding residues only on the 5′ end from 77 nt to 114nt (after adding the 7nt slippery site), and (2) symmetrically expanding both ends from 77 nt to 144 nt (after adding the 7nt slippery site). The asymmetric approach helps determine the shortest FSE length for obtaining the 3_3 pseudoknot (see SHAPE experiment below), and is also realistic for ribosomal interactions. The symmetric expansion helps interpret the full landscape.

For each length, we first extract experimental reactivities for corresponding residues from 144 nt construct, and renormalize them to have the same mean value as the 77 nt construct. Second, we predict the RNA 2D structures using ShapeKnots, along with Gibbs free energies, and then calculate respective Boltzmann probabilities. Third, we sum up probabilities for all structures containing independently folded 3_6 or 3_3 pseudoknot, and display populations in red (3_6) and green (3_3) for each length in Fig. 6. *Although the reactivities used here for different sequence lengths are not the real data from the folded RNA at the given length, we seek to estimate general aspects of the landscape.*

**Figure 6:**
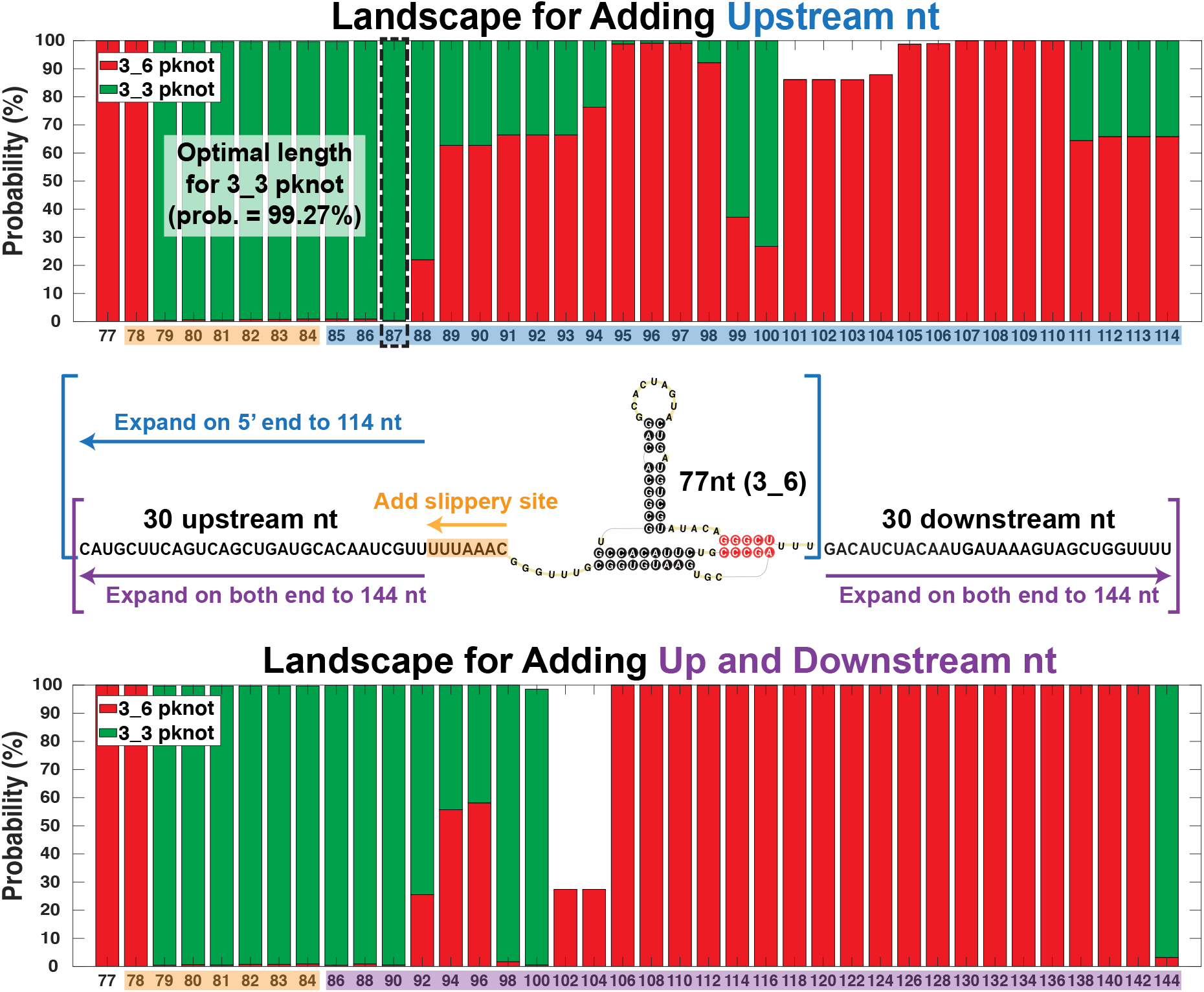
Conformational landscape of the frameshifting element for different sequence lengths predicted by Shape-Knots using reactivities from the 144 nt construct. For each length, probabilities of all structures containing independently folded 3_6 or 3_3 pseudoknots are individually summed. The compositions are colored red (3_6) and green (3_3), respectively. (Top) Landscape for adding upstream nucleotides only to the 77 nt FSE (asymmetric expansion). The optimal sequence length of 87 nt for the 3_3 pseudoknot is in dashed black. (Bottom) Landscape for adding both upstream and downstream nucleotides to the 77 nt FSE (symmetric approach). At 90 nt (87 + 3 downstream nt) the landscape is almost all 3_3.

For the asymmetric expansion (Fig. 6 Top), the 3_6 pseudoknot is dominant for 77 and 78 nt and again around 89-98 nt and 101-114 nt. For other lengths, the 3_3 pseudoknot is dominant, namely over 95% of the landscape for 79-87 nt (same for symmetric expansion with extra downstream nucleotides). In the alternative symmetric expansion (Fig. 6 Bottom), the probability of the 3_6 pseudoknot increases for 92-96 nt, but drops for 98-104 nt. After that, this conformation occupies almost the entire landscape for 106-142 nt. A sudden switch to the dominant 3_3 conformation occurs at 144 nt.

We choose the 87 nt RNA with a probability of 99% 3_3 pseudoknot for further reactivity studies, which indeed yield a dominant 3_3. See Fig. S11. The dominant structure 4_21 is made of the 3_3 pseudoknot and the flanking Stem *S_F_*, with a probability of 99.87%. Moreover, our partition algorithm ^54^ shows that 4_21 is a subgraph of 7_2192 (see Fig. S4), which corresponds to the 144 nt 3_3-pseudoknot-containing structure in Fig. 4B. This indicates that our choice of 87 nt preserves a natural structure adapted by the longer FSE while removing additional flanking nucleotides. The minor structure 4_12, a subgraph of 6_132 (144 nt 3_6-pseudoknot-containing structure), is made of the 3_6 pseudoknot and a 5′ end hairpin.

We also calculate landscapes using NUPACK and ShapeKnots without any SHAPE reactivities (Fig. S12). Many structures emerge, including 3_6, 3_3, 3_5, and 4_7 (the two-pseudoknot fold in Fig. 3 with 3_3 and another pseudoknot at the 3′ end). For NUPACK, the 4_7 pseudoknot instead of 3_3 dominates 79-87 nt, followed by a switch to a dominant 3_6 except at 98-104 nt and 140-144 nt for symmetric expansion. For ShapeKnots, only a small composition of 3_6 is seen; even for 77 nt, we obtain a dominant 3_5 junction. Most of the landscape is occupied by 3_3.

Clearly, the FSE conformation is highly sensitive to length. The confirmation of a 3_3 dominant landscape for 87 nt by SHAPE reactivity data underscores the utility of the above analysis. Flexible Stem 2 may be involved in a switch between the two pseudoknot conformations.

### Comparison with other works

#### Other major and minor FSE conformations in the literature

To relate our three relevant, length-dependent structures for the FSE to recent structural works, we list major and minor FSE structures identified in Table 2. For SARS-CoV FSE, the 3_6 pseudoknot was taken as the consensus structure, ^13,19,58^ and by extension to SARS-CoV-2, it continues to be the prevailing FSE structure. ^15,21,35–37^ Using various techniques and sequence lengths, 12 out of the 18 papers show a major 3_6 pseudoknot: iterative 2D prediction for 68 nt;^23^ 3D modeling and MD simulation for 68 nt;^21^ NMR spectroscopy complemented with DMS footprinting for 68 nt;^35^ 2D, 3D, and MD simulation for 77 nt and 84 nt; ^22^ small-angle X-ray scattering for 85 nt;^15^ DMS-MaPseq for 85 nt;^30^ homology model for 88 nt;^37^ Cryo-EM for 88 nt^29^ and 118 nt;^36^ deep sequencing for 1475 nt.^38^ All except the last two studies use short FSE lengths of 68-88 nt, and most are *in vitro*. As we demonstrated here, short sequences, especially ≤77 nt, tend to have a dominant 6 pseudoknot in the conformational landscape. The remaining 6 papers predict other major FSE conformations instead of the 3_6 pseudoknot: pseudoknot 3_8, or unknotted 2_2 and 2_1.

**Table 2:**
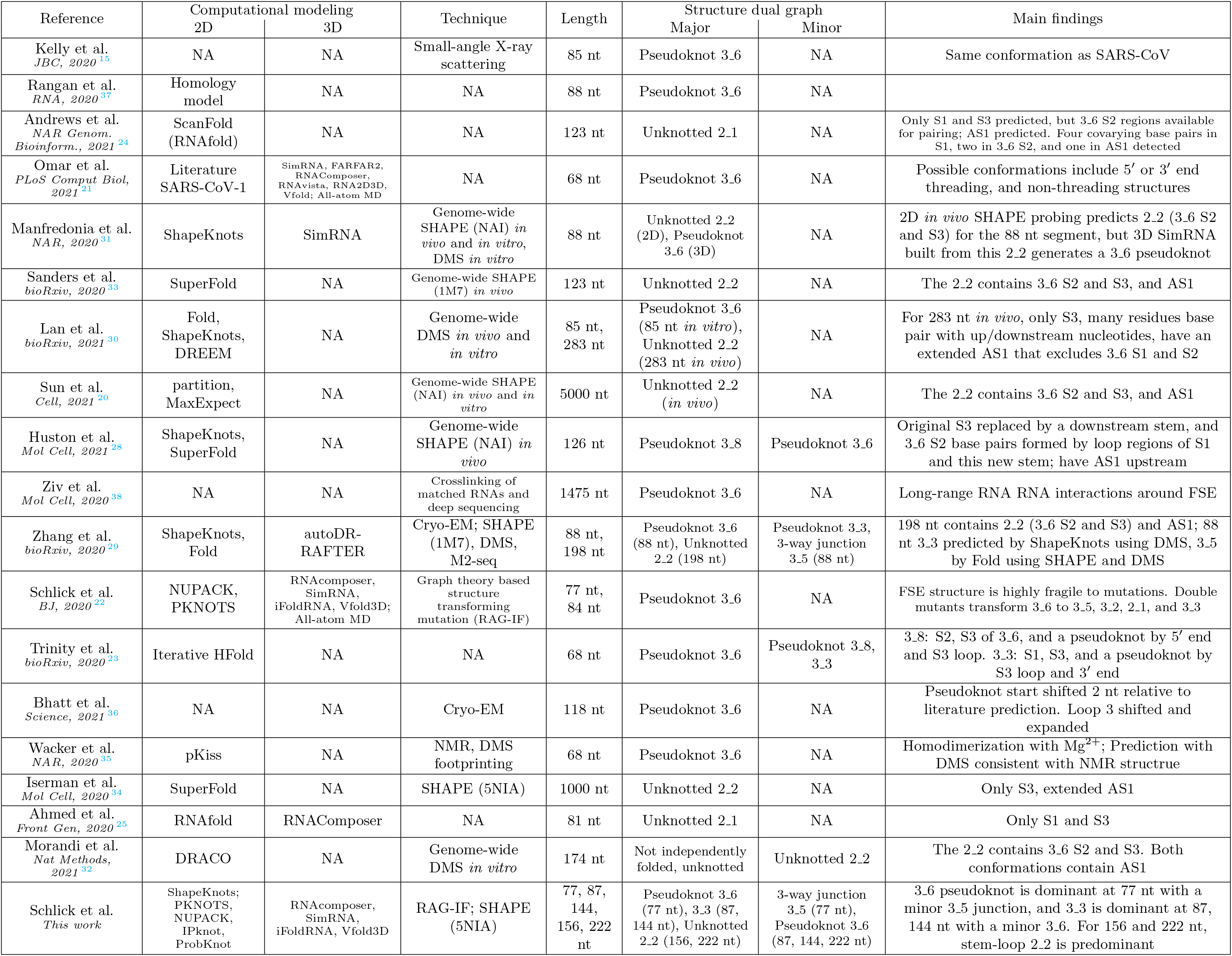
FSE structure prediction in the literature, ordered by date of first archived version of the paper.

The 3_8 kissing hairpin for 126 nt arises in a genome-wide *in vivo* SHAPE experiment paper by Huston et al., ^28^ where parameters for the 3_6 Stem 2 detected by ShapeKnots are hardwired constraints, and Super-Fold ^59^ is applied to predict a consensus structure for the FSE. Their 3_8 pseudoknot (see Fig. S13) replaces the original Stem 3 by a different downstream stem, so that base pairs in the 3_6 Stem 2 involve loop regions of Stem 1 and this new stem.

The 2_2 conformation is a two-stem structure with an internal loop, and it is derived by seven groups by chemical probing long sequences (> 198 nt) extended at the 5′ end. ^20,29–34^ Among these, five groups perform genome-wide probing, four of these *in vivo*. While Stem 3 is conserved in all studies, Stem 2 of 3_6 is predicted by five groups. A common feature of these 2_2 conformations is the replacement of Stem 1 by an upstream (extended) AS1; the exception is the 88 nt structure predicted using genome-wide SHAPE reactivity ^31^ (see Fig. S13). AS1 appears in the 126 nt structure^28^ and in our 2D prediction for 156 nt sequence (Fig. 3), but Stem 1 can co-exist with a shorter AS1.

While this unknotted 2_2 can be partially explained by three groups using 2D prediction programs that do not handle pseudoknots, ^20,33,34^ the authors attribute this to the longer (genome-wide) sequence and the differences caused by *in vivo* vs. *in vitro* experiments. ^30,31^ However, 3D models for these 2_2 conformations do not yield well-defined 3D structures, ^60^ and a 3D structure built from the 88 nt 2_2 actually recovers the 3_6 pseudoknot. ^31^ Combined with its weaker covariation evidence than Stem 1 (Fig. 2, MSA section), this alternative form appears less stable than structures with Stem 1.

To further probe the 2_2 motif, we perform SHAPE and DMS experiments for longer FSE segments of 156 and 222 nt. We find that 2_2 becomes dominant in these two constructs using ShapeKnots prediction (Fig. S14). Moreover, when we compare our chemical probing of different lengths (77, 87, 144, 156, and 222 nt), a sudden increase is observed for both SHAPE and DMS reactivity in the 3′ strand of Stem 1 (residues 13495– 13500) for 156 and 222 nt (Fig. S15). This increase is further supported by the Iserman et al. 1000 nt SHAPE probing, ^34^ which aligns well with our 156 and 222 nt constructs. Hence, a transition from Stem 1 to AS1 might occur between 144 and 156 nt, as residues in the 5′ strand of AS1 are included (all residues of the AS1_5′ strand are included in the 156 nt system).

The 2_1 unknotted conformation contains only Stems 1 and 3 (Fig. S13), so it is a substructure of all our conformers 3_6, 3_3, and 3_5 (Fig. S4). It is dominant in two studies using sequence length 81 nt^25^ and 123 nt.^24^ Both groups use a 2D program which cannot predict pseudoknots, RNAfold. ^61^

Many minor conformations have also been reported as summarized in Table 2 and Fig. S13, including our 3_3 pseudoknot and 3_5 junction captured by Zhang et al., ^29^ who predict the 3_3 pseudoknot by ShapeKnots using DMS data for 88 nt. However, DMS data can only inform about nucleotides A and C, which likely explains why 3_3 is only a minor conformer. Additionally, ShapeKnots is designed for SHAPE reactivities and not DMS. These researchers also obtain 3_5 using both SHAPE and DMS reactivities.

Trinity et al. ^23^ obtain a minor 3_8 and a 3_3 pseudoknot (Fig. S13), but the corresponding 2D structures are different from those we have described. Recall that each graph topology corresponds to multiple 2D structures. In their 3_8 RNA, ^23^ base pairs in Stems 2 and 3 are the same as in our 3_6 in Fig. 1, but the original Stem 1 is replaced by a pseudoknot that binds the 5′ end and the Stem 3 loop. In their 3_3, ^23^ Stem 3 loop and the 3′ end base pair to form the pseudoknot, instead of the 5′ end of the 77 nt FSE and Stem 1 loop.

We also compare chemical probing data for the extended FSE region (residues 13280–13644) in Fig. S16. Our work has generated consistent reactivity profiles for different lengths while the existing data are too heterogeneous to generate consistent models. The *in vivo* genome-wide SHAPE probing in Fig. S16A ^20,28,31^ shows that three groups’ data align poorly with low Pearson correlations (*r* < 0.5), likely due to different reagents and readout technologies. The *in vitro* SHAPE probing including our 222 nt construct in Fig. S16B ^29,31,34^ similarly show poor data agreement and low correlations, except for our 222 nt construct with the Iserman et al. 1000 nt construct (*r* = 0.85, both 5NIA reagent). The *in vitro* DMS probing in Fig. S16C ^29,31,32^ again shows poor data alignment. These comparisons argue for a unified approach as performed in this study.

#### FSE structure and frameshifting efficiency

While clearly the conformational landscape of the FSE is length dependent and fragile to mutations, the relation between structure and frameshifting efficiency is not well understood. In the Cryo-EM study by Bhatt et al., ^36^ the researchers show that mutations of a single residue G13486 in Loop 1 (Fig. 1) reduce frameshifting efficiency, and that deletion of the entire Loop 1 inhibits frameshifting entirely; similarly, removing a single residue A13537 in Loop 3 or the entire Loop 3 reduces frameshifting dramatically (Fig. 7). They suggest that such changes in frameshifting efficiency are caused by altered interactions with the ribosome.

**Figure 7:**
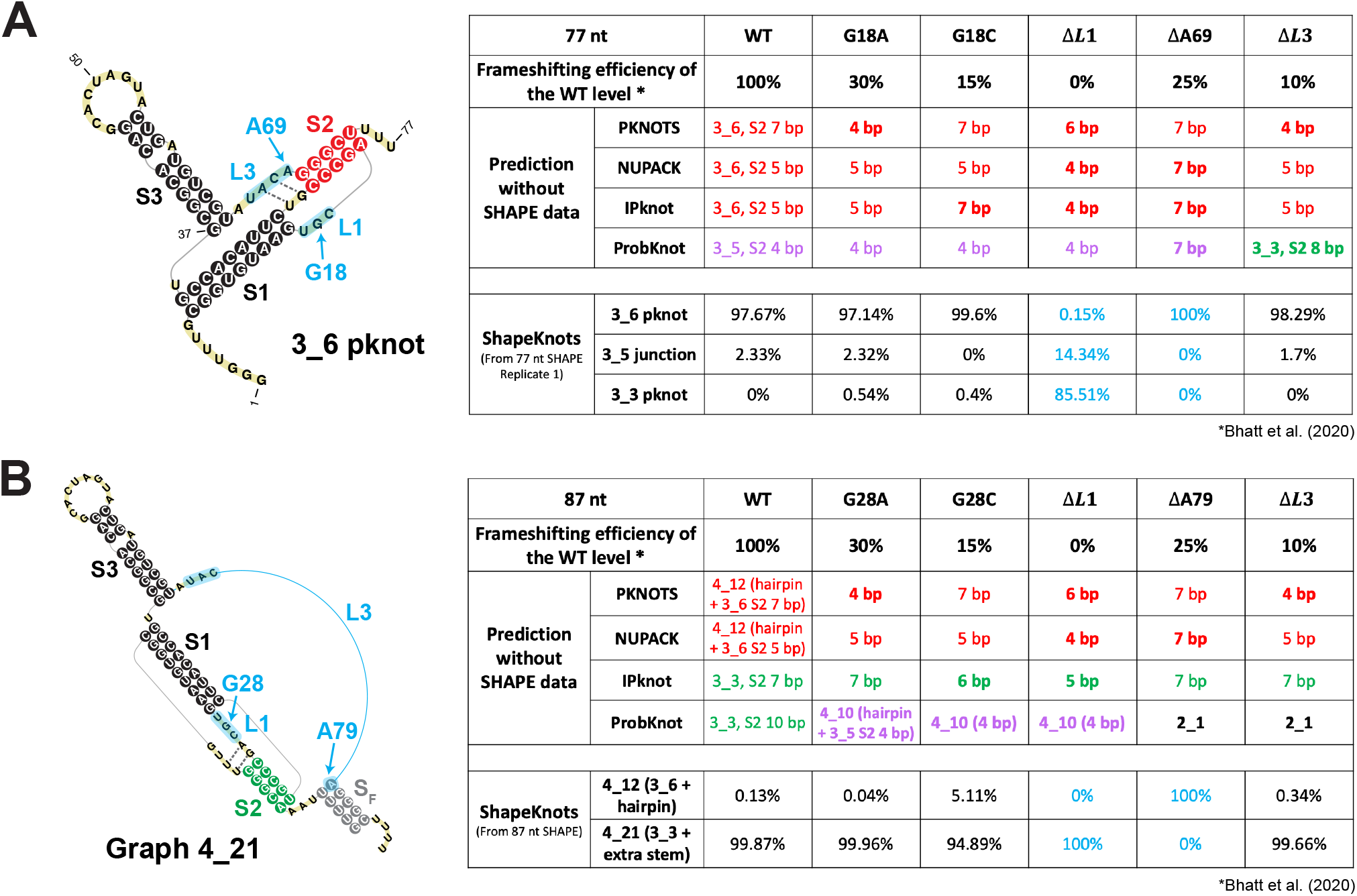
Effects of 5 mutations tested for frameshifting efficiency by Bhatt et al. ^36^ on 2D structure predictions of the 77 nt and 87 nt frameshifting element. (A) (Left) The 77 nt FSE 3_6 pseudoknot with mutation regions labeled in blue. Two weak base pairs for Stem 2 are indicated using dotted lines. (Right) A table showing 2D prediction results for the wildtype and the mutants. The upper half is for 4 programs: 3_6, 3_5, and 3_3 predictions are in red, purple, and green, respectively, with their corresponding Stem 2 lengths (in bold if structure change). The lower half is for ShapeKnots, showing probabilities of 3_6, 3_5, and 3_3. (B) (Left) The 87 nt FSE 2D structure by ShapeKnots, a 4_21 structure made of the 3_3 pseudoknot and a flanking Stem *S_F_*, with mutation regions in blue. (Right) A table showing prediction results using 4 programs and ShapeKnots.

To investigate whether these mutations might also affect the FSE structure and possibly the frameshifting process through a structural change, we predict for each mutation in Fig. 7 the 2D structures of the resulting RNAs for 77 nt and 87 nt FSEs using four prediction programs (PKNOTS, NUPACK, IPknot, and ProbKnot). We also consider predictions with reactivity data for the relevant residues in the original 77 nt and 87 nt FSE constructs. We use red/green/purple consistent with Fig. 1 to highlight resulting 3_6/3_3/3_5 conformations.

Without reactivity data, while the three programs that predict the 3_6 pseudoknot for the wildtype 77 nt FSE continue to predict 3_6 for the mutants, Stem 2’s length is altered. Meanwhile, ProbKnot predicts the 3_5 junction with a 4 bp Stem 2 for the wildtype FSE. This 3_5 Stem 2 is lengthened by deletion of A13537 in Loop 3, but destroyed by deletion of the entire Loop 3, where the alternative 3_3 Stem 2 forms. For 87 nt systems, the 3_3 pseudoknot emerges as dominant structure for the IPknot program for both wildtype and all mutants, consistent with our predictions and SHAPE experiments (Fig. 6 and S11), while only the wildtype for ProbKnot; the structures predicted by ProbKnot for the mutants yield both the 3-way junction and a simple two-stem structure. With SHAPE data, we see that the 3_3 pseudoknot dominates over 3_6 when Loop 1 is deleted, and that all three conformations again play a role in the energy landscape.

The analysis and discussion above underscore the many alternative conformations for the FSE. Even using similar methods such as chemical structure probing can lead to different conformations for different sequence lengths. In particular, the 77 nt FSE 5′ end, which was assumed to be an unpaired spacer region, ^15,21,36^ can form multiple mutually exclusive stems (our 3_3, 3_5 Stem 2, or AS1). The spacer region length is considered to have a critical impact on frameshifting efficiency. ^36^ It is possible that as the elongating ribosome approaches the FSE region, the stems formed with upstream sequence are unwound, and dynamic structural transitions occur among the alternative structures. Moreover, from both our mutant analysis and the 2D structure predictions for the Bhatt et al. mutants, ^36^ we conclude that the FSE structure is highly sensitive to mutations. Altering only a few nucleotides can transform the FSE conformation to an alternative structure or decrease the length of a stem significantly, possibly reducing the frameshifting efficiency.

## Conclusion and discussion: mechanistic implications

Using a combination of graph-based modeling, 2D structure prediction programs, and chemical structure probing data, we have described three alternative structures for the SARS-CoV-2 frameshifting element (Fig. 1). Besides the 3-stem H-type pseudoknot long assumed to be the dominant structure in the literature (3_6 dual graph), ^15,22,35–38^ another 3-stem pseudoknot, HL-type (3_3 dual graph), becomes dominant when 30 nt upstream and 30 nt downstream are added. An unknotted 3-way junction RNA (3_5 dual graph) also emerges as a minor player in the FSE conformational landscape. Using minimal mutations predicted by our genetic algorithm RAG-IF, we can strengthen the prevalence of 3_6, 3_3, and 3_5 in the 77 nt construct using six, three, and two mutations, respectively, highlighting the fragility of the sequence/structure relationship for the FSE. Such motif stabilizing mutations may be useful for anti-viral therapy targeting a specific conformer. The SHAPE reactivity results summarized in Table 1 for all wildtype and mutant replicates confirm our predictions experimentally.

The two main pseudoknot structures are differently intertwined by hydrogen bonding: Stem 1’s loop base pairs with the 3′ end of the FSE to form Stem 2 in the 3_6 pseudoknot, while this loop binds the 5′ end of the FSE in 3_3 (Fig. 1). Both are bulky structures (see three-dimensional views in Fig. 8, Methods, and more details in a separate molecular dynamics paper ^62^). Our estimated conformational landscape as a function of sequence length (Fig. 6) further highlights the plasticity of the FSE, and the likelihood that exogenous factors, such as small molecules, will alter it.

**Figure 8:**
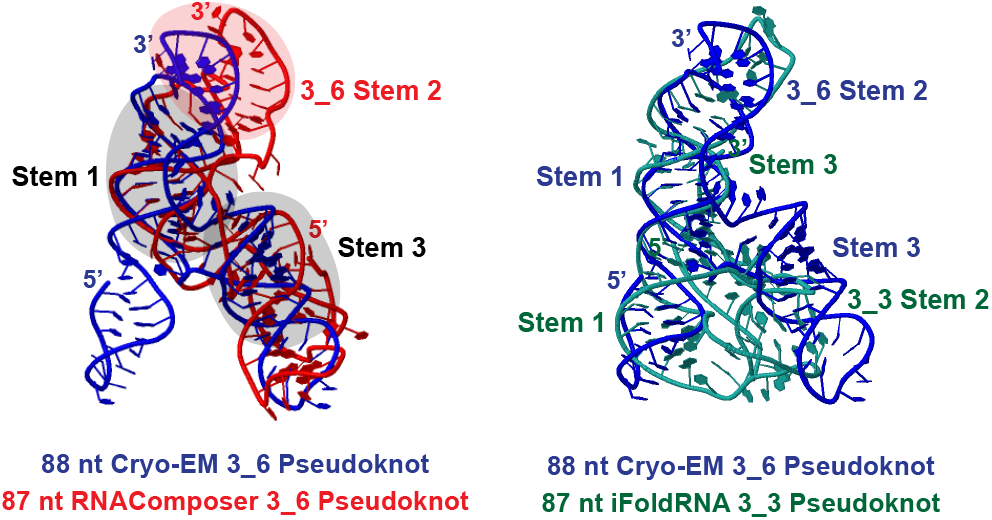
Three-dimensional models of the 87 nt 3_6 and 3_3 pseudoknot. Initial 3D structures are predicted by RNAComposer, ^63^ Vfold3D, ^64^ SimRNA, ^65^ and iFoldRNA, ^66^ and subjected to 1-1.5 *μ*s MD using Gromacs. ^67^ The last 500 ns are used for clustering analysis, and the most populated cluster center by RNAComposer (red)/iFoldRNA (green) is shown here for each system. The 88 nt cryo-EM 3_6 structure derived by Zhang et al. (blue) ^29^ (PDB: 6XRZ) is aligned using Rclick ^68^ for comparison. The three shaded stems of the cryo-EM structure align well with our 3_6 model.

Our multiple sequence alignment of coronaviruses (Fig. 2) similarly underscores the variability of Stem 2 among coronaviruses and high conservation of the FSE sequence and Stem 1. Sequence similarity of the FSE segment (58-98%) is higher than the overall genome similarity (52-89%) among these family members, especially for Sarbecovirus Pangolin-CoV, Bat-CoV, BtRs-BetaCoV, SARS-like WIV1-CoV, SARS-CoV, and BtRf-BetaCoV. The two strands of Stem 1 are highly conserved and a consensus Stem 1 is observed with strong covariation. The flexibility of Stem 2 suggests that this region of the FSE may be involved in a switch between the alternative conformations and/or other biomolecular interactions.

As the sequence length increases beyond 144 nt, additional conformations for the FSE emerge, as Alternative Stem 1 (AS1) ^20,29–34^ becomes more favorable. Our reactivity data coupled with modeling show that a switch between 144 and 156 nt leads to a new peak that corresponds to the 2_2 stem-loop motif with AS1 (Fig. S14). Though so far not associated with a stable 3D structure ^60^ (unlike 3_6, 3_3, and 3_5), this alternative state may be in competition with other FSE forms. Our MSA indicates that this AS1 may be Sarbecovirus-specific, and its covariation evidence is weaker than Stem 1 (Fig. 2). Besides pseudoknots 3_6 and 3_3, three-way junction 3_5, and stem-loop 2_2, alternatives may include two 3_8 pseudoknots, ^23,28^ a different 3_3 pseudoknot (with Stem 2 formed by Stem 3 loop and 3′ end), ^23^ and a two-stem 2_1^24,25^ (Table 2 and Fig. S13). Our work clearly shows that formation of Stem 1 and Alternative Stem 1 are mutually exclusive. Co-transcriptional folding will lead to a preference of AS1, explaining our 2_2 conformation for 156 and 222 nt constructs.

The length-dependent and context-specific conformations for the FSE could be exploited biologically in mechanisms of interactions with the ribosome. The bulky pseudoknot promotes ribosome pausing. ^8–10,69,70^ In the recent 2.3–7 Å resolution Cryo-EM study, ^36^ the researchers observe the 3_6 pseudoknot wedged between the head and body of the small ribosomal subunit.

Besides serving as an obstacle, the FSE may participate more actively in the frameshifting process through conformational transformations.^12–14,71^ Conformational changes might occur during co-transcriptional unfolding, as the elongating ribosome approaches the 5′ strand of AS1 and unwinds it, making the 3′ end of AS1 available for forming Stem 1 and Stem 2 of 3_3 and 3_5. Given estimates for ribosome pausing of ~2.8s between translocations, ^72^ this unwinding of AS1 may promote other conformations and thus conformational transitions. The observations that longer sequences have increased frameshifting suggests that different conformations may indeed be accessible to the frameshifting element.^30^ Once the ribosome moves to the slippery site, the 3_6 pseudoknot may remain as the only viable structure, which may explain the prevalence of 3_6 in many experiments. Different conformations are likely associated with different levels of frameshifting efficiency, and they may be favored differently throughout the virus life cycle to control structural and non-structural protein production. ^71^ Future studies are needed to further discover the role of these alternative conformations in frameshifting and possibly viral packaging.

Similarly, FSE mutations, such as reported recently, ^36^ and proposed in our prior work ^22^ and here, can also affect frameshifting efficiency and FSE structure by helping target specific FSE forms. Drugs that exploit FSE pockets ^15,19,20^ may affect structures, mechanisms, and function. The ribosome anchoring likely affects conformational variability in the realistic context, but the bulkiness of the pseudoknot may be part of the structural signalling as the ribosome unwinds the FSE. We can thus envision at least three avenues for such interference (Fig. 9).

**Figure 9:**
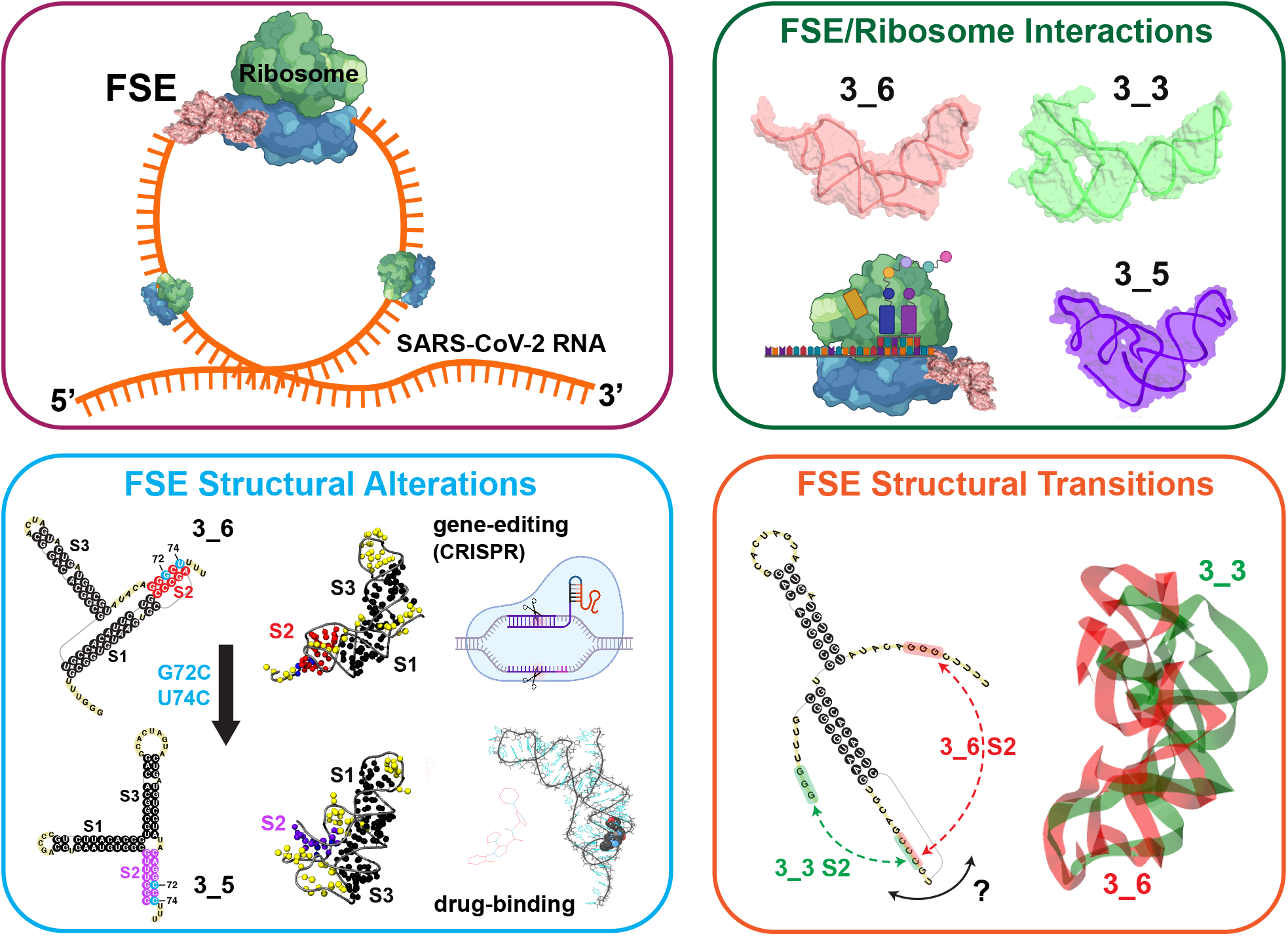
Three avenues for frameshifting interference, with cartoon models for the tertiary systems as modeled by molecular dynamics ^62^ (created with BioRender.com).

### FSE Mutations

Select residues or pockets of the FSE that are vulnerable to mutations by gene editing or drug binding to stabilize a specific FSE conformation or alter the FSE structure and hence interfere with frameshifting. ^15,19,20,22,36,73^

### FSE/Ribosome

Influencing the FSE/ribosome interactions could interfere with the biomolecular recognition process and protein translation occurring in the mRNA entry tunnel. ^36^ For example, Loop 1 or Stem 1 could be good targets here for mutations or drug binding.

### FSE Transitions

Altering conformational rearrangements of the FSE in the heterogeneous landscape could define another avenue. Atomic-level molecular dynamics simulations could help suggest ideas for different inherent motions and threading orientations for the two FSE pseudoknot systems. ^29,62^

As more high resolution structural complexes are reported, it may be possible to hone this picture. Computational studies will clearly be an important part of piecing the clues, as already demonstrated for many aspects of the Covid-19 disease. A combination of coarse-grained modeling as used here for RNA representations with RAG graphs and efficient design of minimal structure-altering mutations, are particularly effective when combined with atomic-level views.

Despite recent suggestions that viral mutations in the spike-protein encoding region may be associated with higher infectivity of recent variants, mutations in the FSE in these variants (Fig. S2) appear random and infrequent, reinforcing this region’s high evolutionary conservation and importance to maintaining viral fitness. In contrast to the need to evolve viral protein inhibitors, the frameshifting inhibitor MTDB was found to be resistant to natural mutations.^45^ The sequence-length and context dependence folding of the FSE and its conformational variability make modeling and experiments infinitely more complicated, but these variations may be a part of the complex machinery that coronaviruses have developed to infect and replicate rapidly and efficiently.

Despite the growing availability of highly effective vaccines against Covid-19, the threat of further variants, coronavirus waves, and other viruses cannot be over-stated. With increased global travel, human invasion of natural forests, and domestication and consumption of wild animal species, more opportunities arise for the jumping of viruses from their natural reservoirs in the animal kingdom to human hosts. A better understanding of the complex structure/function relationship will be critical in this fight against future virus pandemics.

## Methods

### SARS-CoV-2 RNA sequences

We use the official SARS-CoV-2 RNA reference sequence provided by GISAID ^44^ (Accession ID: EPI ISL 402124), 29891 nt. The 84 nt FSE occupies residues 13462–13545, and the spike gene region is residues 21564–25384. Other viral RNA sequences are aligned to the reference by GISAID using *mafft*. ^74^ Specific variants such as the British variant (B.1.1.7 or VUI-202012/01), the South Africa variant (B.1.351 or 501Y.V2), the Brazil variant (P.1), or the New York City variant (B.1.526) were downloaded as fasta files using its search engine.

### Coronavirus FSE multiple sequence alignment and covariation analysis

There are five steps in our coronavirus FSE MSA and covariation analysis:

#### 1. Coronavirus selection

We download 3760 SARS-CoV-2 sequences from GISAID, and 2855 other coronavirus sequences (1129 Alphacoronavirus, 1125 Betacoronavirus, 152 Deltacoronavirus, and 449 Gammacoronavirus) from Virus Pathogen Database and Analysis Resource (ViPR). ^41^ Redundant sequences were removed using CD-HIT ^75^ with similarity threshold 99%, and 1248 sequences remained.

#### 2. Covariance model construction

To build a covariance model, both aligned sequences and a consensus secondary structure are required. Here we input the 222 nt SARS-CoV-2 FSE sequence (residues 13354–13575) and its dominant 2_2 conformation derived by our SHAPE probing (Fig. S14) as the consensus structure into Infernal, ^42^ and run *cmbuild* and *cmcalibrate*.

#### 3. Homologous region identification and alignment

The covariance model built above is used to search for homologous regions in the 1248 coronavirus sequences using Infernal *cmsearch* with option *-A* to output a MSA, and 629 hits are found. We remove duplicates and sequences with unknown characters such as *N*, and 182 sequences remain. Alignment of the 16 top scored sequences with SARS-CoV-2 FSE (Fig. 2) is visualized by Jalview^76^ with a sequence logo generated. The insertions are hidden to save space.

#### 4. Sequence identity calculation

Sequence similarities with SARS-CoV-2 for both the whole genome and the 222 nt FSE segment are calculated using BLAST’s global alignment webserver with default parameters. ^77^

#### 5. Covariation analysis

We input the MSA containing 182 sequences into R-scape with option *-s* and default parameters^43^ to evaluate the 2_2 conformation, and the 3_6, 3_3, and 3_5 structures as well.

### Secondary structure prediction programs

Six 2D structure prediction programs that can handle pseudoknots are used: PKNOTS, ^49^ NUPACK, ^50^ IPknot, ^51^ ProbKnot, ^52^ vsfold5, ^53^ and ShapeKnots. ^56^ Only ShapeKnots can incorporate SHAPE reactivities into the prediction of 2D structures. Except for vsfold5, which works as a webserver, we install the programs locally. Default parameters are used for PKNOTS, NU-PACK *mfe*, ProbKnot, and vsfold5. For IPknot, the parameters are set to level 2 prediction, CONTRAfold scoring model, refinement 1, and base pair weights 2 for level 1 and 16 for level 2. For ShapeKnots, we provide experimental SHAPE reactivity data as input, and calculate all suboptimal structures.

### SHAPE, PairMap

#### Synthesis and purification of *in vitro* transcribed RNA

Various constructs of SARS-CoV-2 FSE (with and without slippery site) were synthesized from Integrated DNA Technologies (IDT). Each construct is flanked by structural RNA adapters ^78^ and a T7 promoter region at the 5′ end. These DNA constructs were used as a template to *in vitro* transcribed RNA using T7 high yield RNA kit (New England Biolabs). The synthesized RNA was DNase treated (TURBODNase), purified using Purelink RNA mini kit (Invitrogen) and quantified with nanodrop.

#### Modification of *in vitro* transcribed RNA

Samples of 6 *μg* of *in vitro* transcribed RNA was denatured at 65 °C for 5 minutes and snap-cooled in ice. After the addition of folding buffer (100 mM KCl, 10 mM MgCl2, 100 mM Bicine, pH 8.3), RNA was incubated at 37 °C for 10 minutes. The folded RNA was treated with either 10 *μl* of Dimethyl sulfate (DMS, 1:10 ethanol diluted) or with 10 *μl* of 5-Nitro Isatoic Anhydride (5NIA, 25 mM final concentration). Subsequently, for negative controls (unmodified RNA) equivalent amount of ethanol and Dimethyl Sulfoxide (DMSO) was added to the folded RNA. The complete reaction mixture was for further incubated for 5 minutes at 37 °C to allow complete modifications of the unpaired RNA nucleotides. DMS treated reaction was quenched using 100 *μl* of 20% beta-mercaptoethanol (*β*-ME). Both the modified and unmodified RNAs were purified using the PurelinkRNA mini kit and quantified with nanodrop.

#### cDNA synthesis, library construction, sequencing, and data processing

Purified RNA from above was reverse transcribed using Gene-specific reverse primer (Table 3) directed against the 3′ RNA adapter sequence and SuperScript II reverse transcriptase under error prone conditions as previously described. ^59^ The resultant cDNA was purified using G50 column (GE healthcare) and subjected to second strand synthesis (NEBNext Second Strand Synthesis Module). For library generation, we designed primers, specific to the 5′ and 3′ RNA adapter sequence (Table 3) and PCR amplified the whole cDNA using the NEB Q5 HotStart polymerase (NEB). Secondary PCR was performed to introduce TrueSeq barcodes. ^59^ All samples were purified using the Ampure XP (Beckman Coulter) beads and Quantification of the libraries was done using Qubit dsDNA HS Assay kit (ThermoFisher). Final libraries were run on Agilent Bioanalyzer for quality check. These TrueSeq libraries were then sequenced as necessary for their desired length, primarily as paired end 2×151 read multiplex runs on MiSeq platform (Illumina). We used the ShapeMapper2 algorithm ^79^ to determine the mutation frequency in both chemically modified (5NIA and DMS treated) and control (DMSO and ethanol treated) RNA samples and to calculate chemical reactivity for each RNA nucleotide using the following equation:

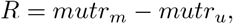

where *R* is the chemical reactivity, *mutr_m_* is the mutation rate calculated for chemically modified RNA and *mutr_u_* is the mutation rate calculated for untreated control RNA samples.

**Table 3:**
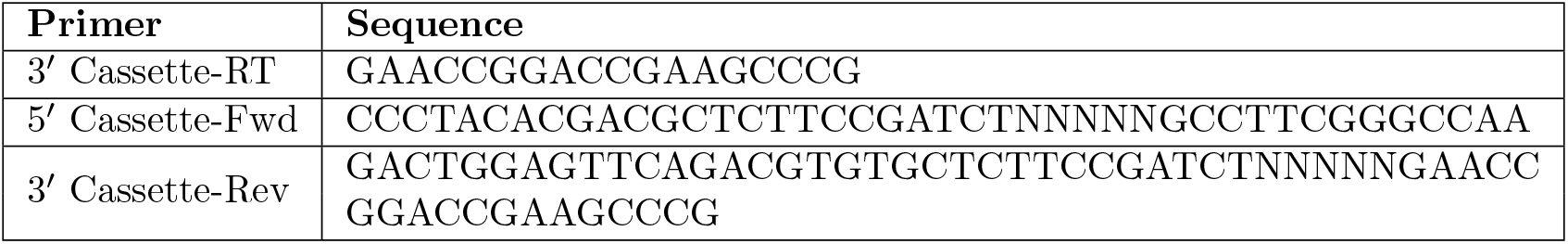
Primers used for the SuperScript II error prone Reverse Transcriptase PCR and library generation

Shapemapper2 was also used to calculate the parse mutations from DMS-MaP sequencing data. ^79^ The resulting parsed mutation files were used in the Pair-MaP pipeline which uses PairMapper and RingMapper to compute and identify correlated mutations in the DMS-MaP sequencing dataset. ^57^ The correlated mutational outputs were plotted with arcPlot. ^57^

SHAPE and DMS reactivities calculated for all the wildtype and mutant constructs are available in the supplementary file *SARS-CoV-2-FSE SNRNASM.xlsx.RAG-IF*

### RAG-IF and mutants design

#### RAG-IF for dual graphs

RAG-IF is an RNA-As-Graphs based inverse folding program that uses genetic algorithm to mutate an RNA sequence so that it folds onto a different target structure (graph) by minimal mutations. It was originally designed and fully automated for tree graphs, ^40^ and modified for dual graphs with manual intervention to select the mutation regions. ^22^ Two prediction programs that can handle pseudoknots are used to determine folding success. Our default options are IPknot and NUPACK (programs *A* and *B*, respectively). However, for 3_6 pseudoknot-strengthening mutant, only ProbKnot predicts a graph (3_5) that is not 3_6 for the wildtype 77 nt. Hence, we substitute default IPknot by ProbKnot in the design of 3_6 PSM. RAG-IF has three steps:

##### 1. Mutation regions and target structure

We identify the smallest mutation region for breaking or forming stems to fold onto the target graph and design a target 2D structure for the target graph.

##### 2. Genetic algorithm

We create an initial population of *N* sequences by randomly assigning nucleotide identities to the mutation regions. Each individual sequence then receives a *fitness* score, which is the number of residues predicted by *Program A* to have the same 2D structure as the target folding, as calculated by the Hamming distance. This population is then subject to *k* iterations of random mutation, crossover, and selection, and those with high fitness are retained as candidates. The algorithm stops once we have enough candidates or the execution time is too long. These candidate sequences are further screened by *Program A* and *B*, and only those that fold onto the target graph by both programs are retained. ^40^

##### 3. Optimization

For each sequence survived above, we remove unnecessary mutations, i.e., the sequence still folds onto the target graph by both prediction programs without these mutations. The remaining mutations are considered minimal. ^40^

#### 3_6 pseudoknot-strengthening mutant

We apply RAG-IF to the 77 nt FSE sequence to predict a 3_6 pseudoknot-strengthening mutant, as illustrated in Fig. S7. ProbKnot predicts a 3_5 junction for the wildtype 77 nt (Fig. S7A). The mutation regions are: residues 1-3 to avoid alternative 3_3 pseudoknot Stem 2, and residues 4 and 67-69 to break the 3_5 Stem 2. For the target 2D structure, we use the 3_6 structure by ShapeKnots (Fig. 4A) with shortened Stem 1. For the genetic algorithm, we create a population of 500 sequences (*N* = 500), and *k* = 500 iterations. The program is terminated when at least 500 candidates are produced or the execution time exceeds 12 hours. RAGIF generates 75 unique sequences with 2-6 mutations. The results are listed in Fig. S7A with illustrative mutations.

To dominate the landscape with the 3_6 conformation (rather than obtain just minimal mutations), we also examine the strength of Stem 2. By screening the 75 mutated sequences by 4 prediction programs (PKNOTS, NUPACK, IPknot, and ProbKnot), we identify the quadruple-mutant [G3U, U4A, C68A, A69C] that all four 2D structure-prediction programs fold onto 3_6 (Fig. S7A). Stem 2 has 7 base pairs using three programs and 5 for ProbKnot. To further strengthen Stem 2, we also mutate residues 18 and 19 in Stem 1 to A, so that they base pair with the UU in the 3′ end. With 6 mutations [G3U, U4A, G18A, C19A, C68A, A69C], all programs predict 9 base pairs for Stem 2.

We test the above 6 mutations on the 144 nt construct. PKNOTS, NUPACK, and IPknot give similar structures as the suboptimal 6_132 structure by Shape-Knots (Fig. 4B): two hairpins in the 5′ end, followed by the 3_6 pseudoknot, and finally a hairpin (highlighted green in Fig. S7B) in the 3′ end. However, ProbKnot predicts a pseudoknot-free structure with only Stems 1 and 3 of 3_6. The 3′ strand of Stem 2 (pink) forms a different stem with the 3′ end. To break this stem and restore 3_6 Stem 2, we add mutation C137A to destroy the middle GC base pair, without altering Stem 2 (pink) and the 3′ end hairpin (green). As expected, the 7-mutant FSE [G40U, U41A, G55A, C56A, C105A, A106C, C137A] yields similar structures containing a 3_6 pseudoknot with 9 base pairs for Stem 2 by all four programs.

#### 3_3 pseudoknot-strengthening mutant

To stabilize the 3_3 pseudoknot, we apply RAG-IF to the 77 nt FSE (Fig. S9) with mutation regions defined by residues 4-6 to form a strengthened 3_3 Stem 2, and residues 70-73 to break the 3_6 Stem 2. We use the same parameters for the genetic algorithm as above. We obtain 20 unique sequences with 1-4 mutations, listed in Fig. S9. After 2D prediction program screening, we consider the triple-mutant [U4C, G71A, G72U] to be the strongest, with 5-7 base pairs for Stem 2 of 3_3.

### 3D models

The 3D structures of the 87 nt FSE were predicted using RNAComposer, ^63^ Vfold3D, ^64^ SimRNA, ^65^ and iFoldRNA. ^66^ One structure from each with the correct graph topology was used for MD simulations.

MD simulations were performed using Gromacs 2020.4, ^67^ with the Amber OL3 forcefield. ^80^ The systems were solvated with TIP3P water molecules in the cubic box whose boundaries extended at least 10 Å from any RNA atom. ^81^ After charge neutralization with randomly placed sodium ions, additional Na^+^ and Cl^−^ ions were added for 0.1 M bulk concentration. The systems were energy minimized via steepest descent and equilibrated with position restraints on the RNA. Simulations were run with a timestep of 2 fs and a SHAKE-like LINCS algorithm ^82^ with constraints on all bonds. The Particle Mesh Ewald method ^83^ was used to treat long-range electrostatics. The equilibration was performed for 100 ps in the NVT ensemble (300 K) and then 100 ps in NPT ensemble (300 K and 1 bar). The RNA and ionic solvent were independently coupled to external heat baths with a relaxation time of 0.1 ps. Production runs were performed for at least 1 *μ*s under NPT, based on when the RMSD stabilized.

Cluster analysis was conducted via Gromos using conformations every 200 ps within the last 500 ns in each simulation using RNA non-H backbone atoms. With a cutoff of 3 Å, the largest cluster occupies 82.3% and 22.4% for 3_6 and 3_3, respectively. The 5′ end in 3_6 is threaded through the ring formed by the 3 stems and extends along the strand of Stem 3 with residues stacked with Stem 3 residues. The 5′ end in 3_3 is not threaded and instead forms a new Stem 2 and pairing with 3′ end. Details of the new MD simulations are described separately in. ^62^

## Supporting information

Chemical Probing Reactivity

## Associated Content

### Supporting Information

SARS-CoV-2 mutation maps; Length effects on 2D predictions; Dual graph partition; Replicate 2 SHAPE analysis for 77 nt and 144 nt FSE; RAG-IF mutant designs; SHAPE reactivity differences; SHAPE replicate alignments; SHAPE analysis for 87 nt FSE; Additional FSE conformational landscape; Other FSE conformations reported in the literature; SHAPE analysis for 156 and 222 nt FSE; Comparisons of chemical probing collected in this study between different FSE lengths; Comparisons of chemical probing by different groups

### SARS-CoV-2-FSE SNRNASM.xlsx

SHAPE and DMS reactivities calculated for all the wild-type and mutant constructs

## Author Information

### Authors

**Qiyao Zhu** – *Courant Institute of Mathematical Sciences, New York University, New York, New York 10012, United States*

**Abhishek Dey** – *Department of Biology, University of North Carolina at Chapel Hill, Chapel Hill, North Carolina 27599, United States*

**Swati Jain** – *Department of Chemistry, New York University, New York, New York 10003, United States*

**Shuting Yan** – *Department of Chemistry, New York University, New York, New York 10003, United States*

### Notes

The authors declare no competing financial interest.

## Acknowledgments

We gratefully acknowledge funding from the National Science Foundation RAPID Award 2030377 from the Divisions of Mathematical Science and of Chemistry, National Institutes of Health R35GM122562 Award from the National Institute of General Medical Sciences, and Philip-Morris International to T. Schlick, and National Institutes of Health grants R35 GM140844, R01 GM101237 and R01 HL111527 to A. Laederach.

## Footnotes

* We use the 3_6, 3_3 pseudoknot and 3_5 junction notations throughout as long as the central FSE region contains these independently folded structures.

## References

(1) Nguyen, T. M.; Zhang, Y.; Pandolfi, P. P. Virus against virus: a potential treatment for 2019-nCov (SARS-CoV-2) and other RNA viruses. Cell Res. 2020, 30, 189–190.

(2) Miao, Z.; Adamiak, R. W.; Antczak, M.; Boniecki, M. J.; Bujnicki, J. M., et al. RNA-Puzzles Round IV: 3D structure predictions of four ribozymes and two aptamers. RNA 2020, 26, 982–995.

(3) Sun, L.-Z.; Zhang, D.; Chen, S.-J. Theory and Modeling of RNA Structure and Interactions with Metal Ions and Small Molecules. Annu. Rev. Biophys. 2017, 46, 227–246.

(4) Brierley, I. Ribosomal frameshifting on viral RNAs. J. Gen. Virol. 1995, 76, 1885–1892.

(5) Atkins, J. F.; Loughran, G.; Bhatt, P. R.; Firth, A. E.; Baranov, P. V. Ribosomal frameshifting and transcriptional slippage: From genetic steganography and cryptography to adventitious use. Nucleic Acids Res. 2016, 44, 7007–7078.

(6) Staple, D. W.; Butcher, S. E. Solution structure of the HIV-1 frameshift inducing stem–loop RNA. Nucleic Acids Res. 2003, 31, 4326–4331.

(7) Brierley, I.; Digard, P.; Inglis, S. Characterization of an efficient coronavirus ribosomal frameshifting signal: requirement for an RNA pseudoknot. Cell 1989, 57, 537–547.

(8) Somogyi, P.; Jenner, A. J.; Brierley, I.; Inglis, S. C. Ribosomal pausing during translation of an RNA pseudoknot. Mol. Cell. Biol. 1993, 13, 6931–6940.

(9) Lopinski, J. D.; Dinman, J. D.; Bruenn, J. A. Kinetics of ribosomal pausing during programmed −1 translational frameshifting. Mol. Cell. Biol. 2000, 20, 1095–1103.

(10) Kim, H. K.; Liu, F.; Fei, J.; Bustamante, C.; Gonzalez, R. L. J.; Tinoco, I. J. A frameshifting stimulatory stem loop destabilizes the hybrid state and impedes ribosomal translocation. Proc. Nat. Acad. Sci., USA 2014, 111, 5538–5543.

(11) Namy, O.; Moran, S.; Stuart, D.; Gilbert, R. J. C.; Brierley, I. A mechanical explanation of RNA pseudoknot function in programmed ribosomal frameshifting. Nature 2006, 441, 244–247.

(12) Ritchie, D. B.; Foster, D. A. N.; Woodside, M. T. Programmed −1 frameshifting efficiency correlates with RNA pseudoknot conformational plasticity, not resistance to mechanical unfolding. Proc. Nat. Acad. Sci., USA 2012, 109, 16167–16172.

(13) Ritchie, D. B.; Soong, J.; Sikkema, W. K. A.; Woodside, M. T. Anti-frameshifting Ligand Reduces the Conformational Plasticity of the SARS Virus Pseudoknot. J. Amer. Chem. Soc. 2014, 136, 2196–2199.

(14) Kelly, J. A.; Woodside, M. T.; Dinman, J. D. Programmed 1 Ribosomal Frameshifting in coronaviruses: A therapeutic target. Virology 2021, 554, 75–82.

(15) Kelly, J. A.; Olson, A. N.; Neupane, K.; Munshi, S.; Emeterio, J. S.; Pollack, L.; Woodside, M. T.; Dinman, J. D. Structural and functional conservation of the programmed 1 ribo-somal frameshift signal of SARS coronavirus 2 (SARS-CoV-2). J. Biol. Chem. 2020, 295, 10741–10748.

(16) Haniff, H. S.; Tong, Y.; Liu, X.; Chen, J. L.; Suresh, B. M.; Andrews, R. J.; Peterson, J. M., et al. Targeting the SARS-CoV-2 RNA Genome with Small Molecule Binders and Ribonuclease Targeting Chimera (RIBOTAC) Degraders. ACS Cent. Sci. 2020, 6, 1713–1721.

(17) Dinman, J. D.; Ruiz-Echevarria, M. J.; Czaplinski, K.; Peltz, S. W. Peptidyl-transferase inhibitors have antiviral properties by altering programmed 1 ribosomal frameshifting efficiencies: Development of model systems. Proc. Nat. Acad. Sci., USA 1997, 94, 6606–6611.

(18) Kinzy, T. G.; Harger, J. W.; Carr-Schmid, A.; Kwon, J.; Shastry, M.; Justice, M.; Dinman, J. D. New Targets for Antivirals: The Ribosomal A-Site and the Factors That Interact with It. Virology 2002, 300, 60–70.

(19) Park, S.-J.; Kim, Y.-G.; Park, H.-J. Identification of RNA Pseudoknot-Binding Ligand That Inhibits the −1 Ribosomal Frameshifting of SARS-Coronavirus by Structure-Based Virtual Screening. J. Amer. Chem. Soc. 2011, 133, 10094–10100.

(20) Sun, L.; Li, P.; Ju, X.; Rao, J.; Huang, W.; Ren, L.; Zhang, S., et al. In vivo structural characterization of the SARS-CoV-2 RNA genome identifies host proteins vulnerable to repurposed drugs. Cell 2021, 184, 1865–1883.e20.

(21) Omar, S. I.; Zhao, M.; Sekar, R. V.; Moghadam, S. A.; Tuszynski, J. A.; Woodside, M. T. Modeling the structure of the frameshift-stimulatory pseudoknot in SARS-CoV-2 reveals multiple possible conformers. PLOS Comput. Biol. 2021, 17, e1008603.

(22) Schlick, T.; Zhu, Q.; Jain, S.; Yan, S. Structure-Altering Mutations of the SARS-CoV-2 Frameshifting RNA Element. Biophys. J. 2021, 120, 1040–1053.

(23) Trinity, L.; Lansing, L.; Jabbari, H.; Stege, U. SARS-CoV-2 ribosomal frameshifting pseudoknot: Improved secondary structure prediction and detection of inter-viral structural similarity. 2020, Article 2020.09.15.298604. bioRxiv. https://doi.org/10.1101/2020.09.15.298604 (accessed September 2020).

(24) Andrews, R. J.; O’Leary, C. A.; Tompkins, V. S.; Peterson, J. M.; Haniff, H. S.; Williams, C.; Disney, M. D.; Moss, W. N. A map of the SARS-CoV-2 RNA structurome. NAR Genom. Bioinform. 2021, 3, lqab043.

(25) Ahmed, F.; Sharma, M.; Al-Ghamdi, A. A.; Al-Yami, S. M.; Al-Salami, A. M., et al. A Comprehensive Analysis of cis-Acting RNA Elements in the SARS-CoV-2 Genome by a Bioinformatics Approach. Front. Genet. 2020, 11, 1385.

(26) Deigan, K. E.; Li, T. W.; Mathews, D. H.; Weeks, K. M.; Tinoco, I. Accurate SHAPE-Directed RNA Structure Determination. Proc. Natl. Acad. Sci. USA 2009, 106, 97–102.

(27) Smola, M. J.; Weeks, K. M. In-cell RNA structure probing with SHAPE-MaP. Nat. Protoc. 2018, 13, 1181–1195.

(28) Huston, N. C.; Wan, H.; Strine, M. S.; de Cesaris Araujo Tavares, R.; Wilen, C. B.; Pyle, A. M. Comprehensive in vivo secondary structure of the SARS-CoV-2 genome reveals novel regulatory motifs and mechanisms. Mol. Cell 2021, 81, 584–598.e5.

(29) Zhang, K.; Zheludev, I. N.; Hagey, R. J.; Wu, M. T.-P.; Haslecker, R., et al. Cryo-electron Microscopy and Exploratory Antisense Targeting of the 28-kDa Frameshift Stimulation Element from the SARS-CoV-2 RNA Genome. 2020, Article 2020.07.18.209270. bioRxiv. https://doi.org/10.1101/2020.07.18.209270 (accessed July 2020).

(30) Lan, T. C. T.; Allan, M. F.; Malsick, L. E.; Khandwala, S.; Nyeo, S. S. Y., et al. Insights into the secondary structural ensembles of the full SARS-CoV-2 RNA genome in infected cells. 2021, Article 2020.06.29.178343. bioRxiv. https://doi.org/10.1101/2020.06.29.178343 (accessed February 2021).

(31) Manfredonia, I.; Nithin, C.; Ponce-Salvatierra, A.; Ghosh, P.; Wirecki, T. K., et al. Genome-wide mapping of SARS-CoV-2 RNA structures identifies therapeutically-relevant elements. Nucleic Acids Res. 2020, 48, 12436–12452.

(32) Morandi, E.; Manfredonia, I.; Simon, L. M.; Anselmi, F., et al. Genome-scale deconvolution of RNA structure ensembles. Nat. Methods 2021, 18, 249–252.

(33) Sanders, W.; Fritch, E. J.; Madden, E. A.; Graham, R. L.; Vincent, H. A.; Heise, M. T.; Baric, R. S.; Moorman, N. J. Comparative analysis of coronavirus genomic RNA structure reveals conservation in SARS-like coronaviruses. 2020, Article 2020.06.15.153197. bioRxiv. https://doi.org/10.1101/2020.06.15.153197 (accessed June 2020).

(34) Iserman, C.; Roden, C. A.; Boerneke, M. A.; Sealfon, R. S.; McLaughlin, G. A.; Jungreis, I.; Fritch, E. J., et al. Genomic RNA Elements Drive Phase Separation of the SARS-CoV-2 Nucleocapsid. Mol. Cell 2020, 80, 1078–1091.

(35) Wacker, A.; Weigand, J. E.; Akabayov, S. R.; Altincekic, N.; Bains, J. K., et al. Secondary structure determination of conserved SARS-CoV-2 RNA elements by NMR spectroscopy. Nucleic Acids Res. 2020, 48, 12415–12435.

(36) Bhatt, P. R.; Scaiola, A.; Loughran, G.; Leibundgut, M.; Kratzel, A., et al. Structural basis of ribosomal frameshifting during translation of the SARS-CoV-2 RNA genome. Science 2021, eabf3546.

(37) Rangan, R.; Zheludev, I. N.; Hagey, R. J.; Pham, E. A.; Wayment-Steele, H. K.; Glenn, J. S.; Das, R. RNA genome conservation and secondary structure in SARS-CoV-2 and SARS-related viruses: a first look. RNA 2020, 26, 937–959.

(38) Ziv, O.; Price, J.; Shalamova, L.; Kamenova, T.; Goodfellow, I.; Weber, F.; Miska, E. A. The Short- and Long-Range RNA-RNA Interactome of SARS-CoV-2. Mol. Cell 2020, 80, 1067–1077.e5.

(39) Chen, S. J. Graph, pseudoknot, and SARS-CoV-2 genomic RNA: A biophysical synthesis. Biophys. J. 2021, 120, 980–982.

(40) Jain, S.; Tao, Y.; Schlick, T. Inverse Folding with RNA-As-Graphs Produces a Large Pool of Candidate Sequences with Target Topologies. J. Struct. Biol. 2020, 209, 107438.

(41) Pickett, B. E.; Sadat, E. L.; Zhang, Y.; Noronha, J. M.; Squires, R. B.; Hunt, V.; Liu, M., et al. ViPR: an open bioinformatics database and analysis resource for virology research. Nucleic Acids Res. 2012, 40, D593–598.

(42) Nawrocki, E. P.; Eddy, S. R. Infernal 1.1: 100-fold faster RNA homology searches. Bioinformatics 2013, 29, 2933–2935.

(43) Rivas, E.; Clements, J.; Eddy, S. R. A statistical test for conserved RNA structure shows lack of evidence for structure in lncRNAs. Nat. Methods 2017, 14, 45–48.

(44) Elbe, S.; Buckland-Merrett, G. Data, disease and diplomacy: GISAID’s innovative contribution to global health. Glob. Chall. 2017, 1, 33–46.

(45) Neupane, K.; Munshi, S.; Zhao, M.; Ritchie, D. B.; Ileperuma, S. M.; Woodside, M. T. Anti-frameshifting ligand active against SARS Coronavirus-2 is resistant to natural mutations of the frameshift-stimulatory pseudoknot. J. Mol. Biol. 2020, 432, 5843–5847.

(46) Tavares, R. d. C. A.; Mahadeshwar, G.; Wan, H.; Huston, N. C.; Pyle, A. M. The global and local distribution of RNA structure throughout the SARS-CoV-2 genome. J. Virol. 2020, 95, e02190–20.

(47) Lange, S. J.; Maticzka, D.; Möhl, M.; Gagnon, J. N.; Brown, C. M.; Backofen, R. Global or local? Predicting secondary structure and accessibility in mRNAs. Nucleic Acids Res. 2012, 40, 5215–5226.

(48) Andrews, R. J.; Roche, J.; Moss, W. N. ScanFold: an approach for genome-wide discovery of local RNA structural elements—applications to Zika virus and HIV. PeerJ 2018, 6, e6136.

(49) Rivas, E.; Eddy, S. R. A dynamic programming algorithm for RNA structure prediction including pseudoknots. J. Mol. Biol. 1999, 285, 2053–2068.

(50) Dirks, R. M.; Pierce, N. A. A partition function algorithm for nucleic acid secondary structure including pseudoknots. J. Comput. Chem 2003, 24, 1664–1677.

(51) Sato, K.; Kato, Y.; Hamada, M.; Akutsu, T.; Asai, K. IPknot: fast and accurate prediction of RNA secondary structures with pseudoknots using integer programming. Bioinform. 2011, 27, i85–i93.

(52) Bellaousov, S.; Mathews, D. H. ProbKnot: fast prediction of RNA secondary structure including pseudoknots. RNA 2010, 16, 1870–1880.

(53) Dawson, W. K.; Fujiwara, K.; Kawai, G. Prediction of RNA Pseudoknots Using Heuristic Modeling with Mapping and Sequential Folding. PLOS ONE 2007, 2, 1–7.

(54) Petingi, L.; Schlick, T. Partitioning and Classification of RNA Secondary Structures into Pseudonotted and Pseudoknot-free Regions Using a Graph-Theoretical Approach. IAENG Int. J. Comput. Sci. 2017, 44, 241–246.

(55) Siegfried, N. A.; Busan, S.; Rice, G. M.; Nelson, J. A. E.; Weeks, K. M. RNA motif discovery by SHAPE and mutational profiling (SHAPE-MaP). Nat. Methods 2014, 11, 959–965.

(56) Hajdin, C. E.; Bellaousov, S.; Huggins, W.; Leonard, C. W.; Mathews, D. H.; Weeks, K. M. Accurate SHAPE-directed RNA secondary structure modeling, including pseudoknots. Proc. Natl. Acad. Sci. USA 2013, 110, 5498–5503.

(57) Mustoe, A. M.; Lama, N. N.; Irving, P. S.; Olson, S. W.; Weeks, K. M. RNA base-pairing complexity in living cells visualized by correlated chemical probing. Proc. Natl. Acad. Sci. USA 2019, 116, 24574–24582.

(58) Plant, E. P.; Pérez-Alvarado, G. C.; Jacobs, J. L.; Mukhopadhyay, B.; Hennig, M.; Dinman, J. D. A three-stemmed mRNA Pseudoknot in the SARS Coronavirus Frameshift Signal. PLOS Biol. 2005, 3, e172.

(59) Smola, M. J.; Rice, G. M.; Busan, S.; Siegfried, N. A.; Weeks, K. M. Selective 2′-hydroxyl acylation analyzed by primer extension and mutational profiling (SHAPE-MaP) for direct, versatile and accurate RNA structure analysis. Nat. Protoc. 2015, 10, 1643–1669.

(60) Rangan, R.; Watkins, A. M.; Chacon, J.; Kladwang, W.; Zheludev, I. N., et al. De novo 3D models of SARS-CoV-2 RNA elements from consensus experimental secondary structures. Nucleic Acids Res. 2021, 49, 3092–3108.

(61) Lorenz, R.; Bernhart, S. H.; Siederdissen, C. H.; Tafer, H.; Flamm, C.; Stadler, P. F.; Hofacker, I. L. ViennaRNA Package 2.0. Algorithms Mol. Biol. 2011, 6, 26.

(62) Yan, S.; Jain, S.; Zhu, Q.; Schlick, T. Length dependent 3D structures and motions of SARS-CoV-2 frameshifting pseudoknot and alternative pseudoknot elements. 2021, in preparation.

(63) Biesiada, M.; Purzycka, K. J.; Szachniuk, M.; Blazewicz, J.; Adamiak, R. W. Automated RNA 3D Structure Prediction with RNAComposer. Methods Mol. Biol. 2016, 1490, 199–215.

(64) Xu, X.; Chen, S.-J. Hierarchical Assembly of RNA Three-Dimensional Structures Based on Loop Templates. J. Phys. Chem. 2018, 122, 5327–5335.

(65) Boniecki, M. J.; Lach, G.; Dawson, W. K.; Tomala, K.; Lukasz, P., et al. SimRNA: a coarse-grained method for RNA folding simulations and 3D structure prediction. Nucleic Acids Res. 2016, 44, e63–e63.

(66) Krokhotin, A.; Houlihan, K.; Dokholyan, N. V. iFoldRNA v2: folding RNA with constraints. Bioinform. 2015, 31, 2891–2893.

(67) Abraham, M. J.; Murtola, T.; Schulz, R., et al. GROMACS: High performance molecular simulations through multi-level parallelism from laptops to supercomputers. SoftwareX 2015, 1-2, 19–25.

(68) Nguyen, M. N.; Verma, C. Rclick: A web server for comparison of RNA 3D structures. Bioinformatics 2015, 31, 966–968.

(69) Chen, J.; Petrov, A.; Johansson, M.; Tsai, A.; O’Leary, S. E.; Puglisi, J. D. Dynamic pathways of −1 translational frameshifting. Nature 2014, 512, 328–332.

(70) Caliskan, N.; Katunin, V. I.; Belardinelli, R.; Peske, F.; Rodnina, M. V. Programmed –1 Frameshifting by Kinetic Partitioning during Impeded Translocation. Cell 2014, 157, 1619–1631.

(71) Moomau, C.; Musalgaonkar, S.; Khan, Y. A.; Jones, J. E.; Dinman, J. D. Structural and Functional Characterization of Programmed Ribosomal Frameshift Signals in West Nile Virus Strains Reveals High Structural Plasticity Among cis-Acting RNA Elements. J. Biol. Chem. 2016, 291, 15788–15795.

(72) Wen, J.; Lancaster, L.; Hodges, C.; Zeri, A.; Yoshimura, S. H.; Noller, H. F.; Bustamante, C.; Tinoco, I. Following translation by single ribosomes one codon at a time. Nature 2008, 452, 598–603.

(73) Schlick, T. In silico drug discovery targeting the SARS-CoV-2 frameshifting RNA element. Joint Mathematics Meetings: AMS Special Session on Advances in Computational Biomedicine II, Virtual, January 8, 2021.

(74) Katoh, K.; Misawa, K.; Kuma, K.; Miyata, T. MAFFT: a novel method for rapid multiple sequence alignment based on fast Fourier transform. Nucleic Acids Res. 2002, 30, 3059–3066.

(75) Fu, L.; Niu, B.; Zhu, Z.; Wu, S.; Li, W. CD-HIT: accelerated for clustering the next generation sequencing data. Bioinformatics 2012, 28, 3150–3152.

(76) Clamp, M.; Cuff, J.; Searle, S. M.; Barton, G. J. The Jalview Java alignment editor. Bioinform. 2004, 20, 426–427.

(77) Altschul, S. F.; Gish, W.; Miller, W.; Myers, E. W.; Lipman, D. J. Basic local alignment search tool. J. Mol. Biol. 1990, 215, 403–410.

(78) Wilkinson, K. A.; Merino, E. J.; Weeks, K. M. Selective 2′-hydroxyl acylation analyzed by primer extension (SHAPE): Quantitative RNA structure analysis at single nucleotide resolution. Nat. Protoc. 2006, 1, 1610–1616.

(79) Busan, S.; Weeks, K. M. Accurate detection of chemical modifications in RNA by mutational profiling (MaP) with ShapeMapper 2. RNA 2018, 24, 143–148.

(80) Zgarbová, M.; Otyepka, M.; Šponer, J.; Mládek, A.; Banáš, P.; Cheatham, T. E.; Jurečka, P. Refinement of the Cornell et al. Nucleic Acids Force Field Based on Reference Quantum Chemical Calculations of Glycosidic Torsion Profiles. J. Chem. 2011, 7, 2886–2902.

(81) Jorgensen, W. L.; Chandrasekhar, J.; Madura, J. D.; Impey, R. W.; Klein, M. L. Comparison of simple potential functions for simulating liquid water. J. Chem. Phys. 1983, 79, 926–935.

(82) Hess, B.; Bekker, H.; Berendsen, H. J. C.; Fraaije, J. G. E. M. LINCS: A linear constraint solver for molecular simulations. J. Comput. Chem. 1997, 18, 1463–1472.

(83) Essmann, U.; Perera, L.; Berkowitz, M. L.; Darden, T.; Lee, H.; Pedersen, L. G. A smooth particle mesh Ewald method. J. Chem. Phys. 1995, 103, 8577–8593.

