## Supplementary Information for "To knot or not to knot: Multiple conformations of the SARS-CoV-2 frameshifting RNA element"

Tamar Schlick,*,†,‡,§ Qiyao Zhu,‡ Abhishek Dey,¶ Swati Jain,† Shuting Yan,† and Alain Laederach*,¶

†Department of Chemistry, 100 Washington Square East, Silver Building, New York University, New York, NY 10003 U.S.A.

‡Courant Institute of Mathematical Sciences, New York University, 251 Mercer St., New York, NY 10012 U.S.A.

¶Department of Biology, University of North Carolina at Chapel Hill, Chapel Hill, NC 27599

§NYU-ECNU Center for Computational Chemistry, NYU Shanghai, Shanghai 200062, P.R. China

#### SARS-CoV-2 mutation maps

GISAID[1] uses *mafft*[2] to align all other viral sequences to the reference sequence. We downloaded the alignment prepared on February 12, 2021, and then produced the mutation maps for the FSE and the spike gene segment by counting mutations in the aligned sequences for every residue in the segment (Fig. S1). The maximum mutation count is 5541 in the FSE region, and 436160 in the spike region. However, most of the FSE residues have mutation counts $\leq 10$, so we choose a scale of 0 to 200 for the FSE mutation map to show enough details for these residues. For the spike region, we choose a scale of 0 to 2000, and we see similar overall bar heights, suggesting an order of magnitude more mutations than the FSE region.

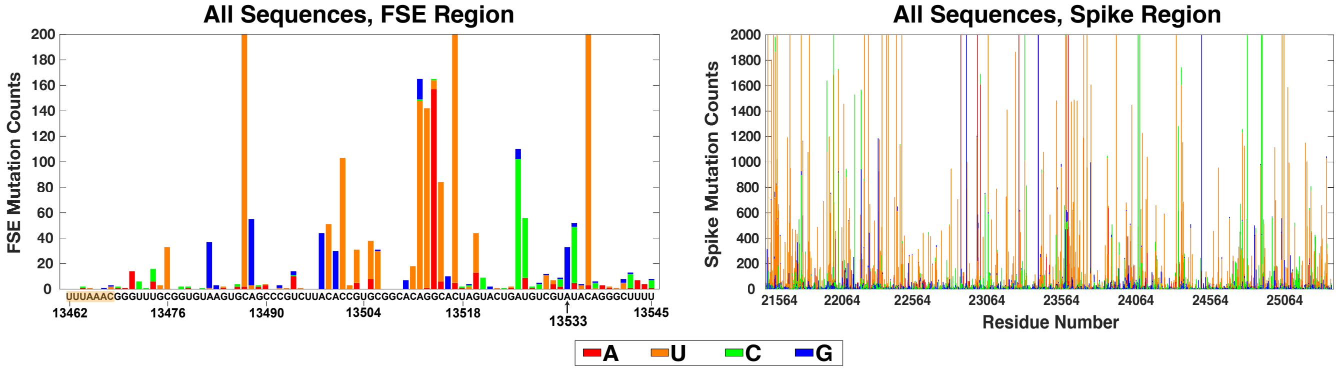


Figure S1: SARS-CoV-2 RNA mutation maps as available on GISAID on February 12, 2021. (Left) All sequence mutation map (459421 viral sequences) for the 84 nt FSE region, with mutations colored based on nucleotide identity. Mutations are counted for the reference sequence of 29891 nt. (Right) Mutation map for the 3822 nt spike region.

For each of the British, South Africa, Brazil, and New York City variant, we downloaded the variant sequences from GISAID and then aligned them with the reference following similar steps as GISAID's MSA:

1. Align each sequence to the reference by the following command line:

```
mafft --thread -1 [input] > [output]
```

2. Separate the sequences into two groups: Group 1 for sequences that bring insertions into the reference when performing Step 1 (*G1*), and Group 2 for the others (*G2*).

3. Group 1 sequences are aligned together with the reference using a gap opening penalty $-10$:

```
mafft --retree 3 --maxiterate 10 --thread -1 --nomemsave --op 10 [G1] > [G1_aligned]
```

4. Add group 2 sequences into the alignment:

```
mafft --thread 1 --nomemsave --keeplength --add [G2] [G1_aligned] > [msa]
```

The variant mutation map (Fig. S2) is then produced using the final MSA *msa* by the same protocol as the all sequence map.

The recent highly transmissible British variant (B.1.1.7) is associated with 6 amino acid substitutions and 2 deletions in the spike protein. Among the 3575 British variant sequences, all of them have 5-11 mutations in the spike gene, while only 4 sequences have mutations (all single nucleotide) in the FSE segment. The mutation maps are shown in Fig. S2A. We find that 7 residues in the spike segment have mutation rates $> 99.7\%$, and 6 of them correspond to the 6 amino acid substitutions. For the FSE segment, only 3 residues are mutated: C13506U twice, C13517U once, and A13533G once.

We conduct the same variant analysis for the recent concerning South Africa variant (B.1.351) as well, and we see similar results. Among the 898 South Africa variant sequences, all of them have 4-12 mutations in the spike gene, while only 5 have mutations (all single nucleotide) in the FSE segment. In the mutation maps (Fig. S2B), 7 residues in the spike segment have mutation rates $> 87.9\%$. In the FSE, 3 residues are mutated once, and the mutation C13517U seen above occurs twice.

Among the 94 Brazil variant (P.1) sequences, all of them have 9-13 mutations in the spike gene, while no mutation is seen in the FSE region (Fig. S2C). Moreover, 12 residues in the spike segment have mutation rates $> 94.6\%$.

Finally, among the 707 New York City variant (B.1.526) sequences, all of them have 2-9 mutations in the spike gene, while only 4 have mutations (all single nucleotide) in the FSE segment. In the mutation maps (Fig. S2D), 4 residues in the spike segment have mutation rates $> 96.3\%$. In the FSE, we see two new mutations: C13511U three times and C13536U once.

#### Folding predictions for 4 pseudoknot-containing RNAs

To examine how sequence lengths affect 2D structure prediction programs, we selected four experimentally solved RNAs with pseudoknots from the Protein Data Bank (PDB) https://www.rcsb.org. The RNAs are: lysine riboswitch (PDB ID: 3DIO), lariat-capping ribozyme (6GYV), glmS ribozyme (2GCV), and T-Box riboregulator (6UFG). The 3D structures are extracted from respective PDB files, and corresponding 2D structures are determined using the RNApdbee webserver.[3]

Pseudoknot substructures in these RNAs are identified and taken to define "short" systems for 2D structure prediction. An equal number of nucleotides are added to both ends of the lysine riboswitch and lariat-capping ribozyme pseudoknots to reach 120 nt. Only the $3'$ ends of the glmS ribozyme and T-Box riboregulator pseudoknot are expanded to 120 nt, because no upstream nucleotides are relevant. The five programs PKNOTS, NUPACK, IPknot, ProbKnot, and vsfold5 are used to predict 2D structures for three sizes ("short", 120 nt, and all). The predictions are compared to the 2D structures extracted from the experimental structures to assess the number of residues that are correctly base paired. The accuracies are calculated as the percentage of correctly base-paired residues in the total number of residues (Fig. S3).

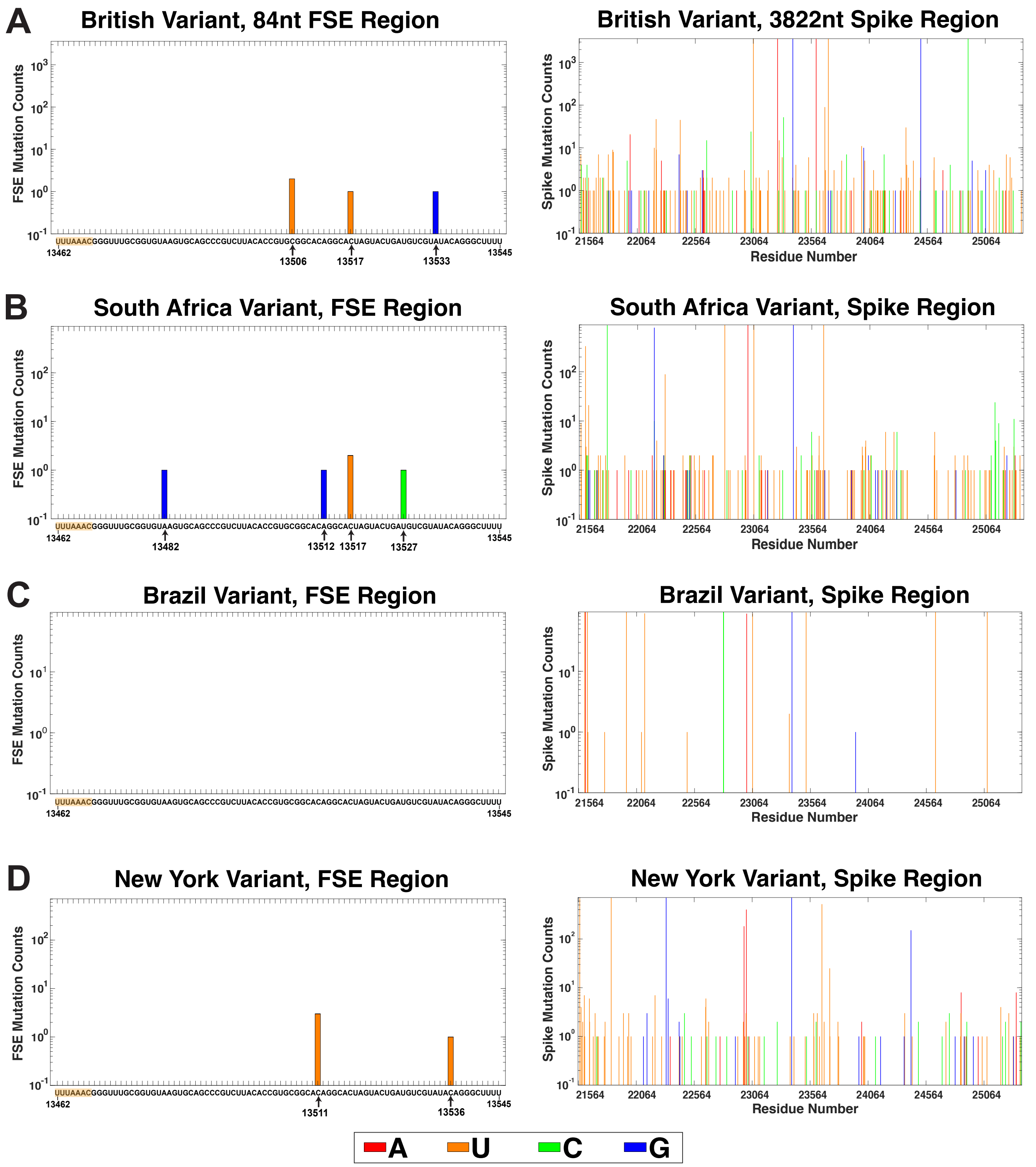


Figure S2: SARS-CoV-2 RNA mutation maps for the (A) British (B.1.1.7) variant, (B) South Africa (B.1.351) variant, (C) Brazil (P.1) variant, and (D) New York (B.1.526) variant for (left) the 84 nt FSE region and (right) the spike region.

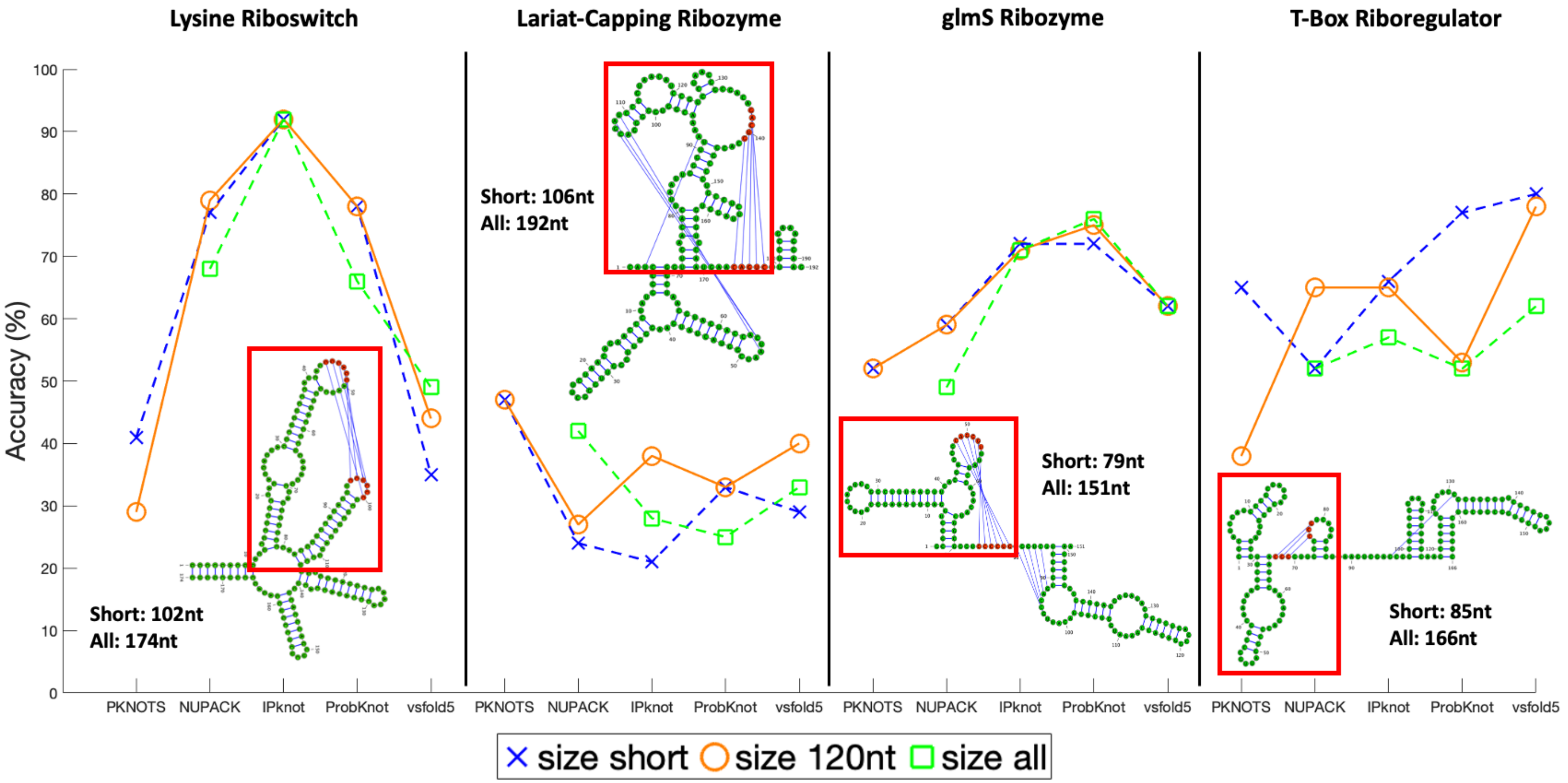


Figure S3: Length effects on 2D structure predictions for four RNAs with pseudoknots. For each RNA, its experimentally determined 2D structure is shown, with the pseudoknot substructure boxed in red. Three length scales are taken for predictions using five programs: "short" for the pseudoknot substructure, 120 nt created by adding equal number of nucleotides on both sides of the pseudoknot, and "all" from the whole RNA structure. For the four RNAs as listed left to right, the pseudoknot lengths are 102, 106, 79, and 85 nt, respectively, and the total lengths are 174, 192, 151, and 166 nt, respectively. Prediction accuracies with respect to the experimental structures are determined by the relative number of correct base pairs for the three lengths, using PDB structures 3DIO for lysine riboswitch, 6GYV for lariat-capping ribozyme, 2GCV for glmS ribozyme, and 6UFG for T-box riboregulator. Although the accuracy varies with the method and RNA, the 120 nt window recommended in the literature appears reasonable in general.

### Dual graph partition and substructure

We developed a partition algorithm that divides a dual graph into its component subgraphs while keeping pseudoknots and junction intact.[4] Here, we apply it to several important dual graphs mentioned in this paper (Fig. S4).

Dual graph 3_6 and 3_5 cannot be divided further, but graph 3_3 can be divided into 2_3, which represents the pseudoknot formed by intertwining Stems 1 and 2, and 2_1 representing two hairpins Stems 1 and 3. The two subgraphs are combined by overlapping one common vertex (Stem 1).

Dual graph 4_7 represents the two-pseudoknot structure predicted by NUPACK for the 84 nt FSE (Fig. 3), and it can be divided into 3_3 and 2_3. The 2_3 graph represents the extra pseudoknot formed by the loop region of Stem 3 and the 3′ end.

Dual graph 4_12 represents the 3_6-containing structure predicted for the 87 nt FSE by ShapeKnots (Fig. S11), and here the 3_6 subgraph is detected by our partition algorithm. Meanwhile, this 4_12 is a subgraph of 6_132, which represents the 144 nt minor 3_6-containing structure (Fig. 4).

Similarly, dual graph 4_21 represents the 87 nt 3_3-containing structure (Fig. S11), with a flanking Stem $S_F$. Though 4_21 cannot be partitioned further, it is identified as a subgraph of 7_2192, which represents the 144 nt major 3_3-containing structure (Fig. 4).

Although graphs 3_6, 3_5, and 4_21 cannot be partitioned further, we can still see clearly the substructures contained in them. The vertices (Stems 1 and 3) and edges in graph 2_1 (omitting self-loops) are contained in 3_6, 3_3, and 3_5. Likewise, graph 3_3 is contained in 4_21.

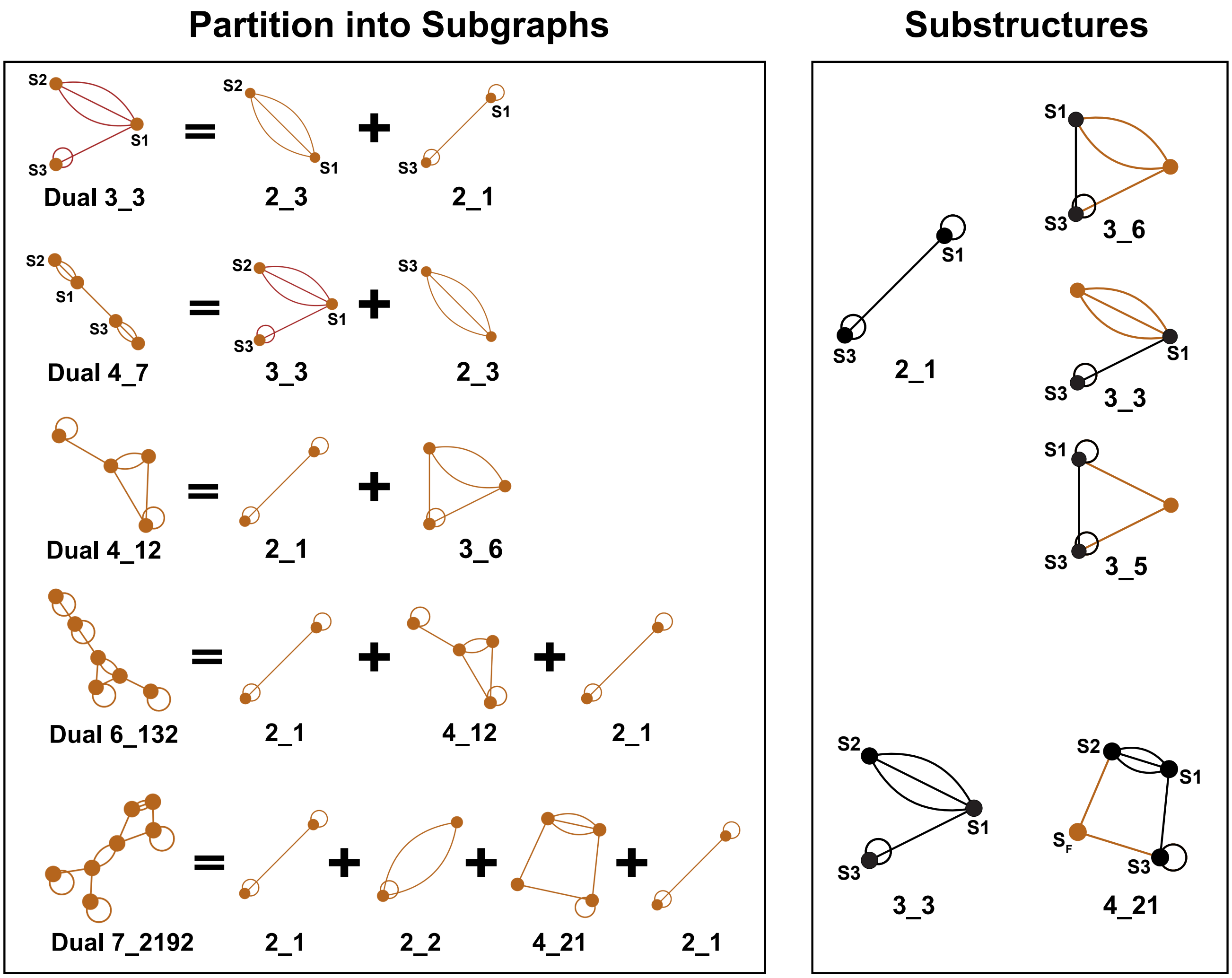


Figure S4: Dual graph partition and substructure.

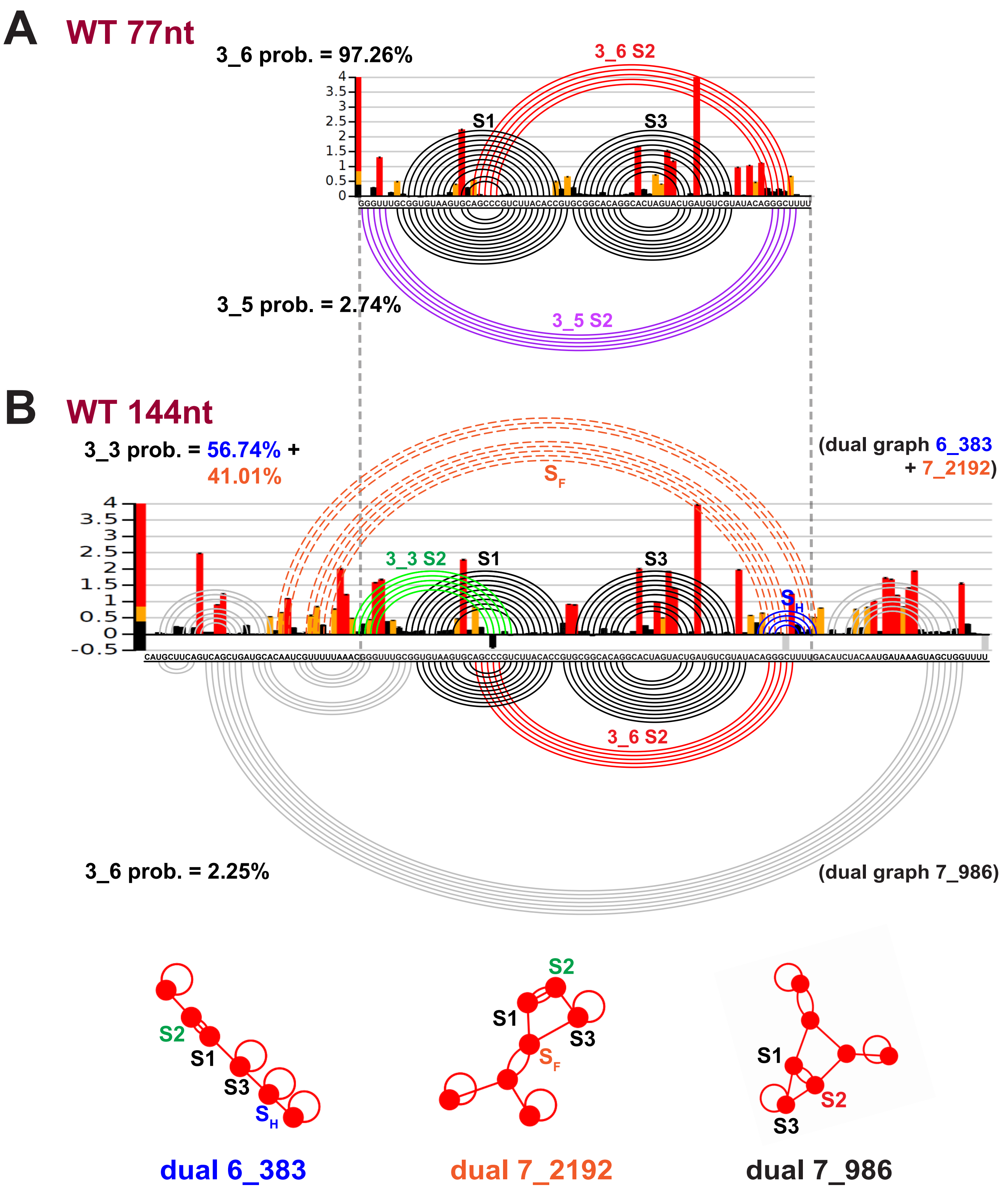


Figure S5: Replicate 2 SHAPE reactivity analysis for SARS-CoV-2 frameshifting element for 77 nt and 144 nt. (A) The SHAPE reactivity for the 77 nt construct is plotted by bars, with red/yellow/black representing high/medium/low reactivity. The arc plot at top shows the dominant 3_6 pseudoknot predicted by ShapeKnots, and at bottom is the minor 3_5. (B) (Top) SHAPE reactivity and ShapeKnots predictions for 144 nt construct. The arc plot at top shows two major 3_3-containing structures predicted by ShapeKnots, with common base pairs except two flanking stems (orange) replaced by a downstream hairpin $S_H$ (blue), and at bottom is the minor 3_6-containing structure. (Bottom) Labeled dual graphs for the three structures predicted for 144 nt.

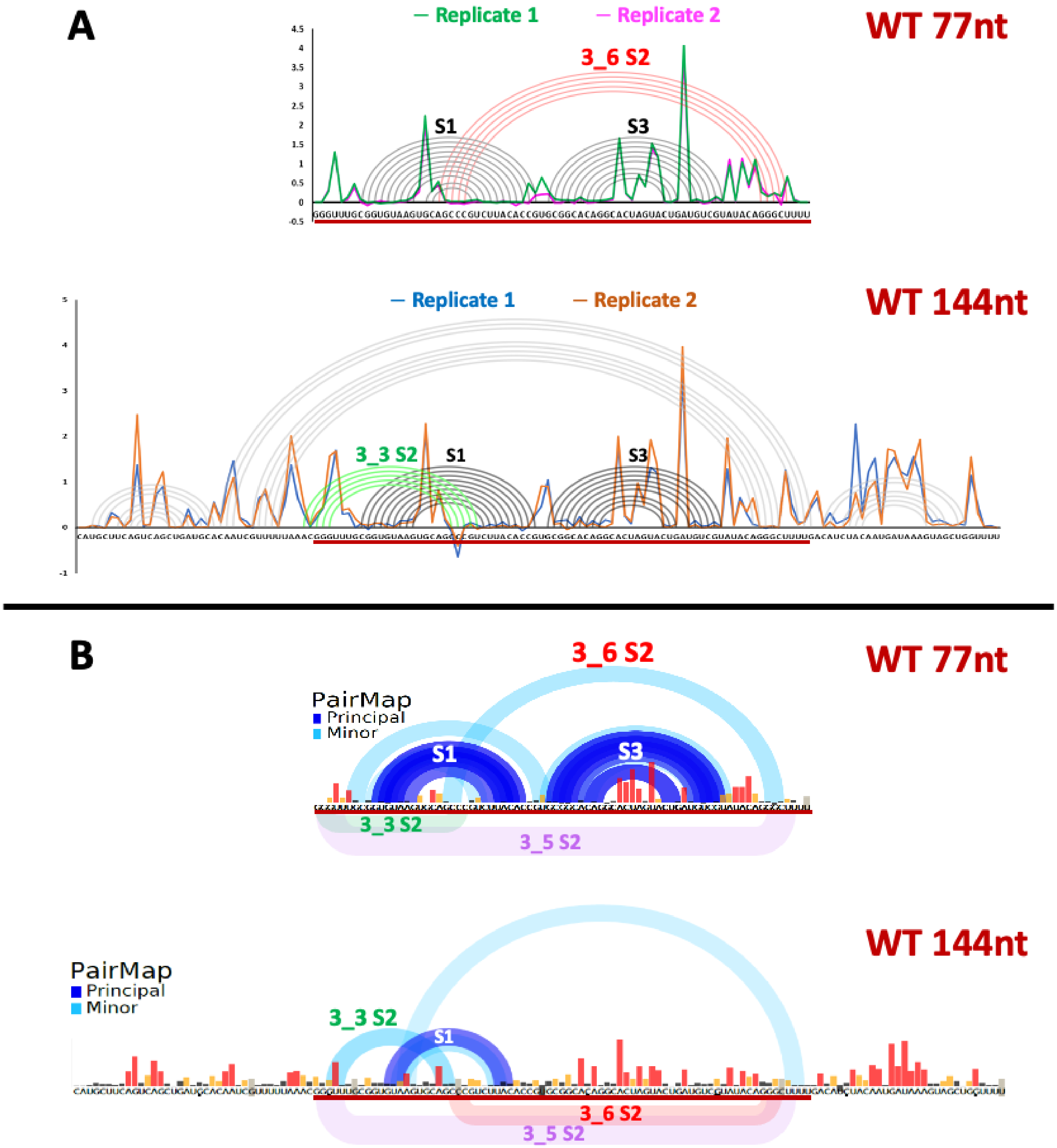


Figure S6: Replicate comparative analysis for the SHAPE reactivity data and application for wildtype 77 nt and 144 nt. (A) The reactivity data for the (top) two 77 nt and (bottom) two 144 nt replicates are aligned for comparison. The consensus dominant structures by ShapeKnots are superimposed as arc plots. (B) PairMap analysis for (top) 77 nt and (bottom) 144 nt construct based on DMS structure probing of the RNA followed by correlated mutation analysis. This analysis reveals regions of nucleotides that are likely to form pairing interactions, but absence of correlated mutations does not indicate lack of pairing.

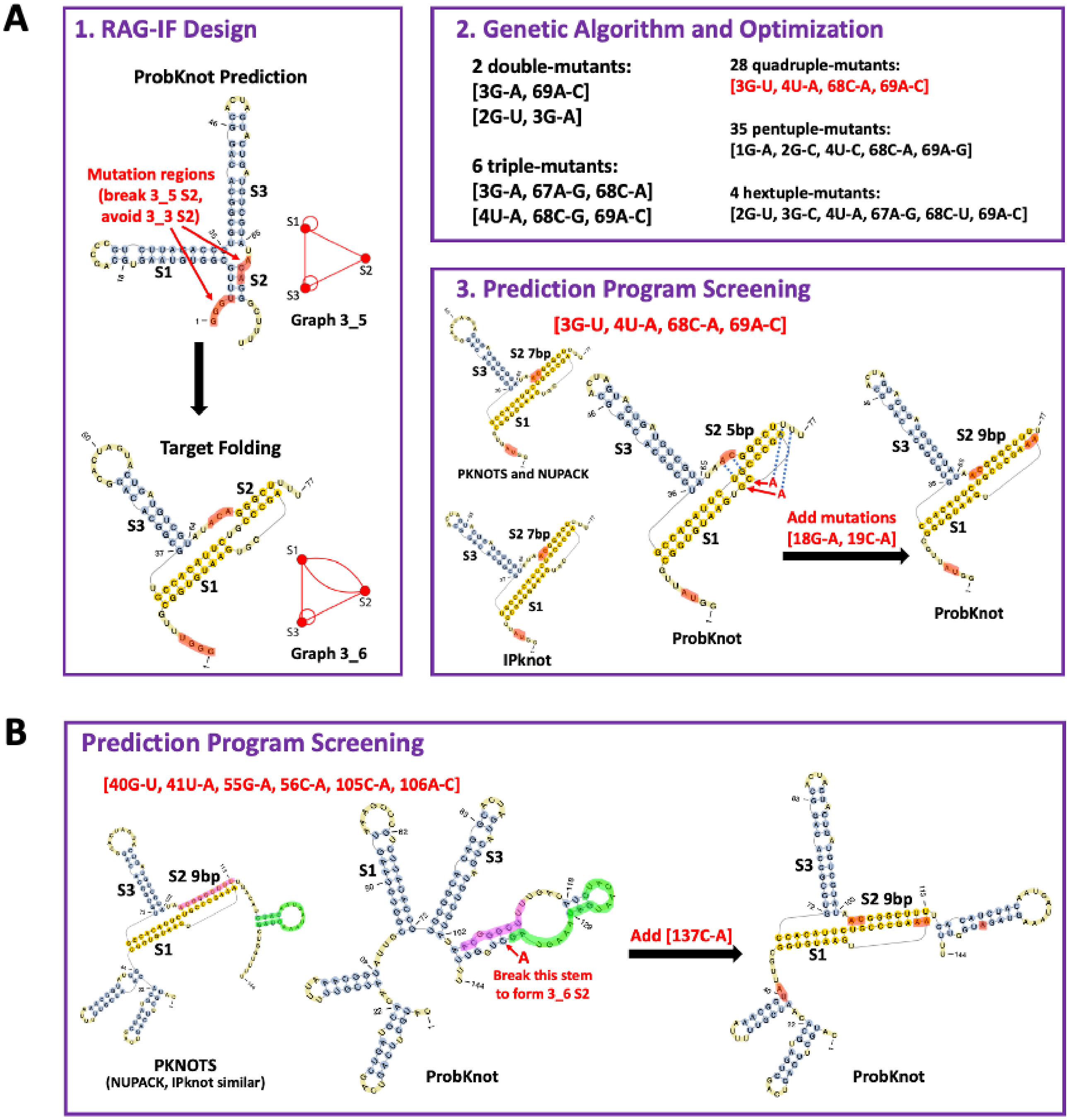


Figure S7: Design of the 3_6 pseudoknot-strengthening mutants (PSMs). (A) 3_6 PSM for 77 nt. Panel 1 shows the mutation regions and the target folding for RAG-IF. Panel 2 lists minimal mutation results. Panel 3 shows 2D prediction program screening for the strongest mutant [3G-U, 4U-A, 68C-A, 69A-C], and identification of two additional mutations [18G-A, 19C-A] based on ProbKnot. (B) 3_6 PSM for the 144 nt construct. Program screening using 6 programs identify another mutation based on ProbKnot. See Methods of text for details.

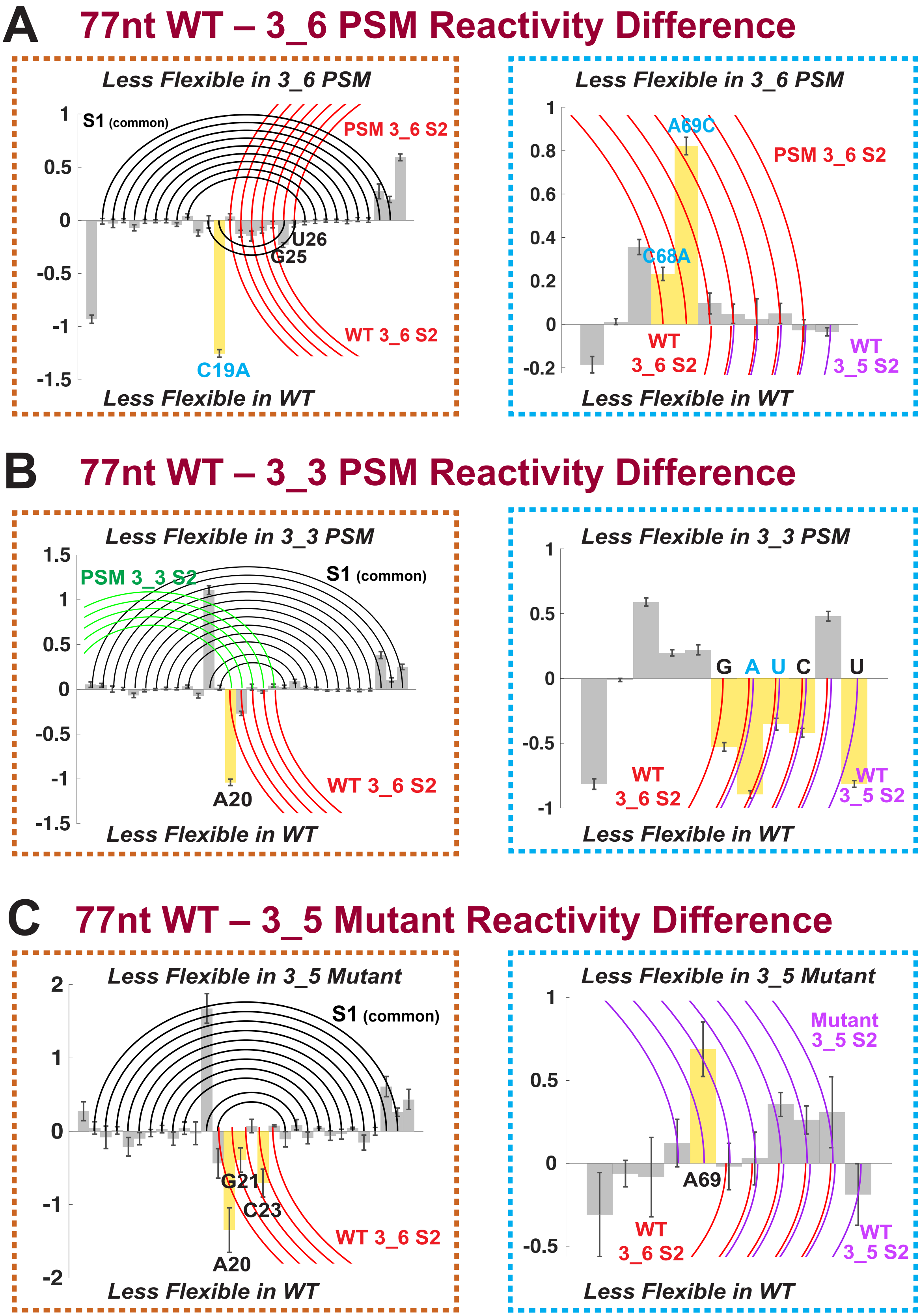


Figure S8: Reactivity differences for two key regions between the 77 nt wildtype and (A) 3_6 PSM, (B) 3_3 PSM, or (C) 3_5 Mutant. Positive/negative differences indicating lower flexibility in the mutant/wildtype. Base pairs in the mutant are plotted by arcs at top, and base pairs in the wildtype at bottom. Critical residues for reactivity comparisons are highlighted.

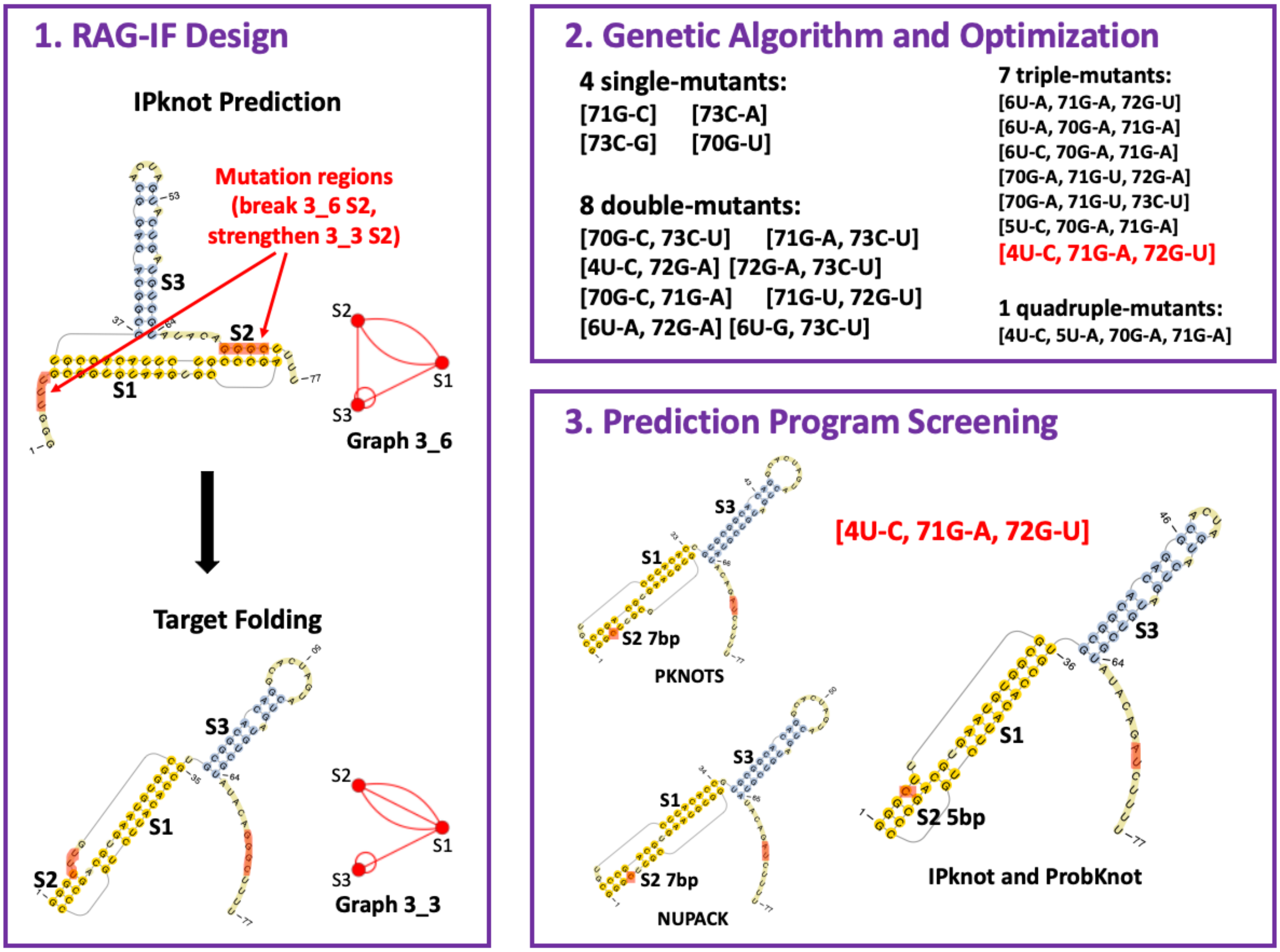


Figure S9: Design of the 3_3 pseudoknot-strengthening mutant (PSM). Panel 1 shows the mutation regions and the target folding for RAG-IF. Panel 2 lists minimal mutation results. Panel 3 shows 2D prediction program screening for the strongest mutant [4U-C, 71G-A, 72G-U].

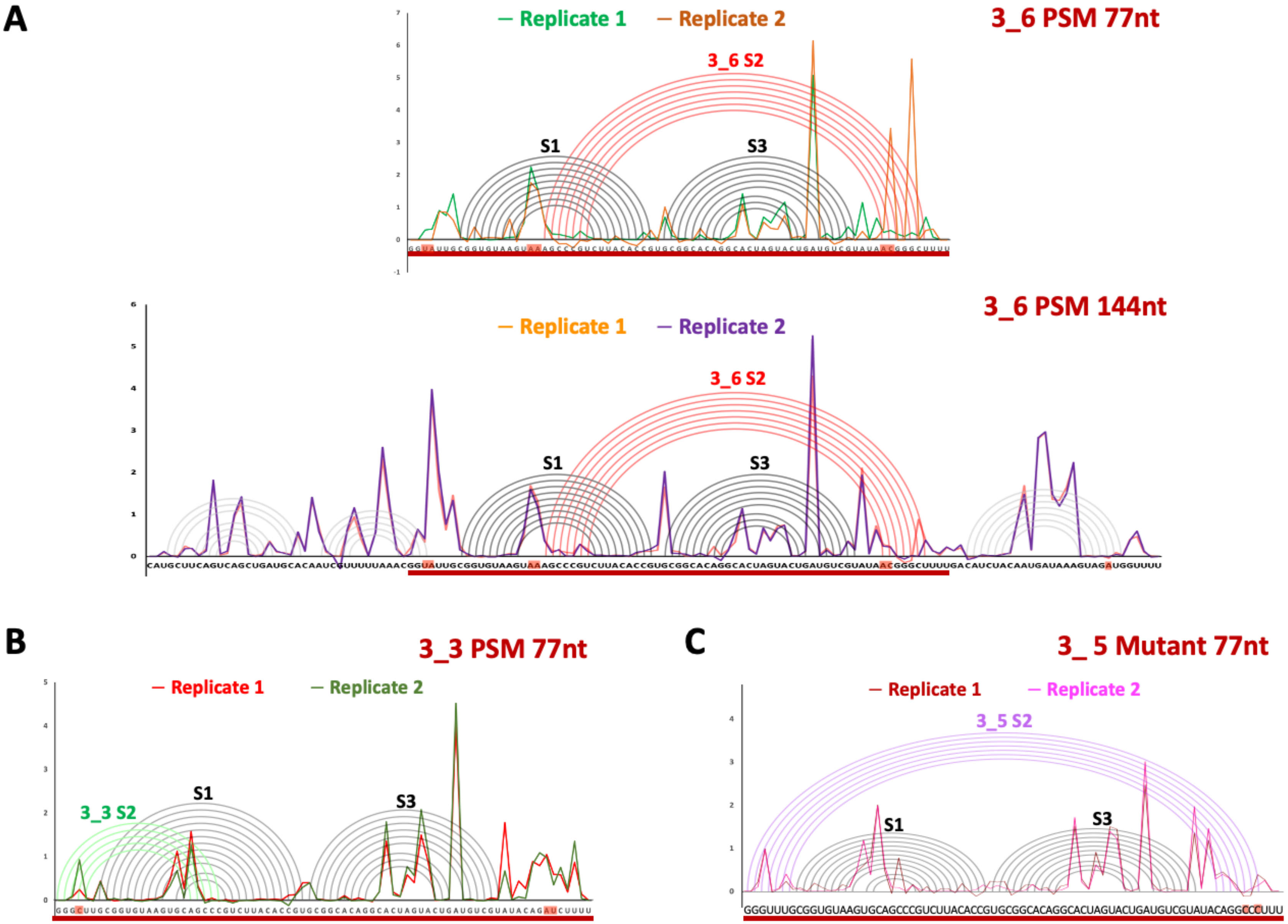


Figure S10: Alignments of mutant replicates and their consensus structures for (A) 3_6 PSM 77 nt and 144 nt, (B) 3_3 PSM 77 nt, and (C) 3_5 Mutant 77 nt.

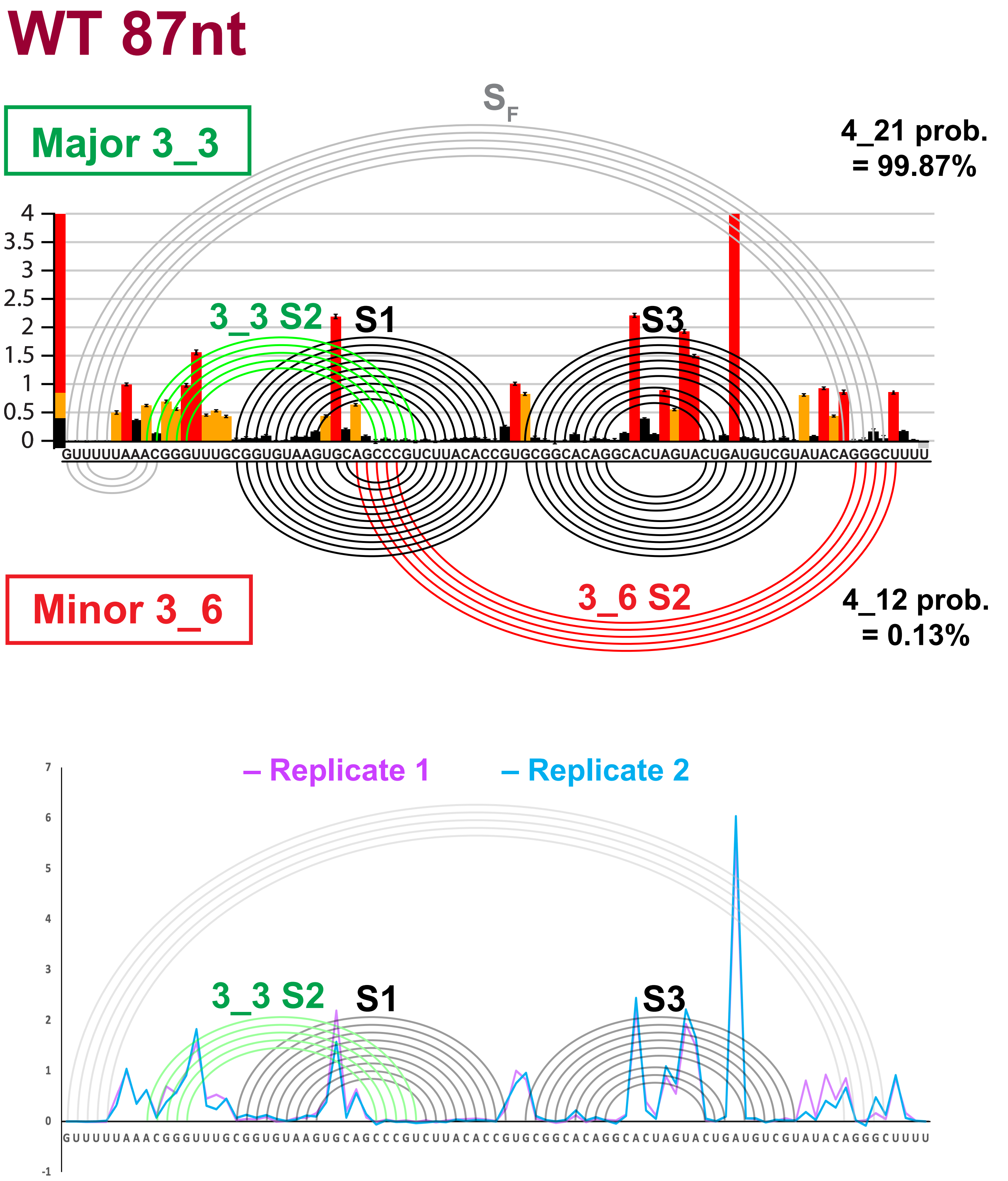


Figure S11: SHAPE reactivity analysis for wildtype 87 nt frameshifting element. (Top) SHAPE reactivity and ShapeKnots predictions for Replicate 1. (Bottom) Alignment of two replicates.

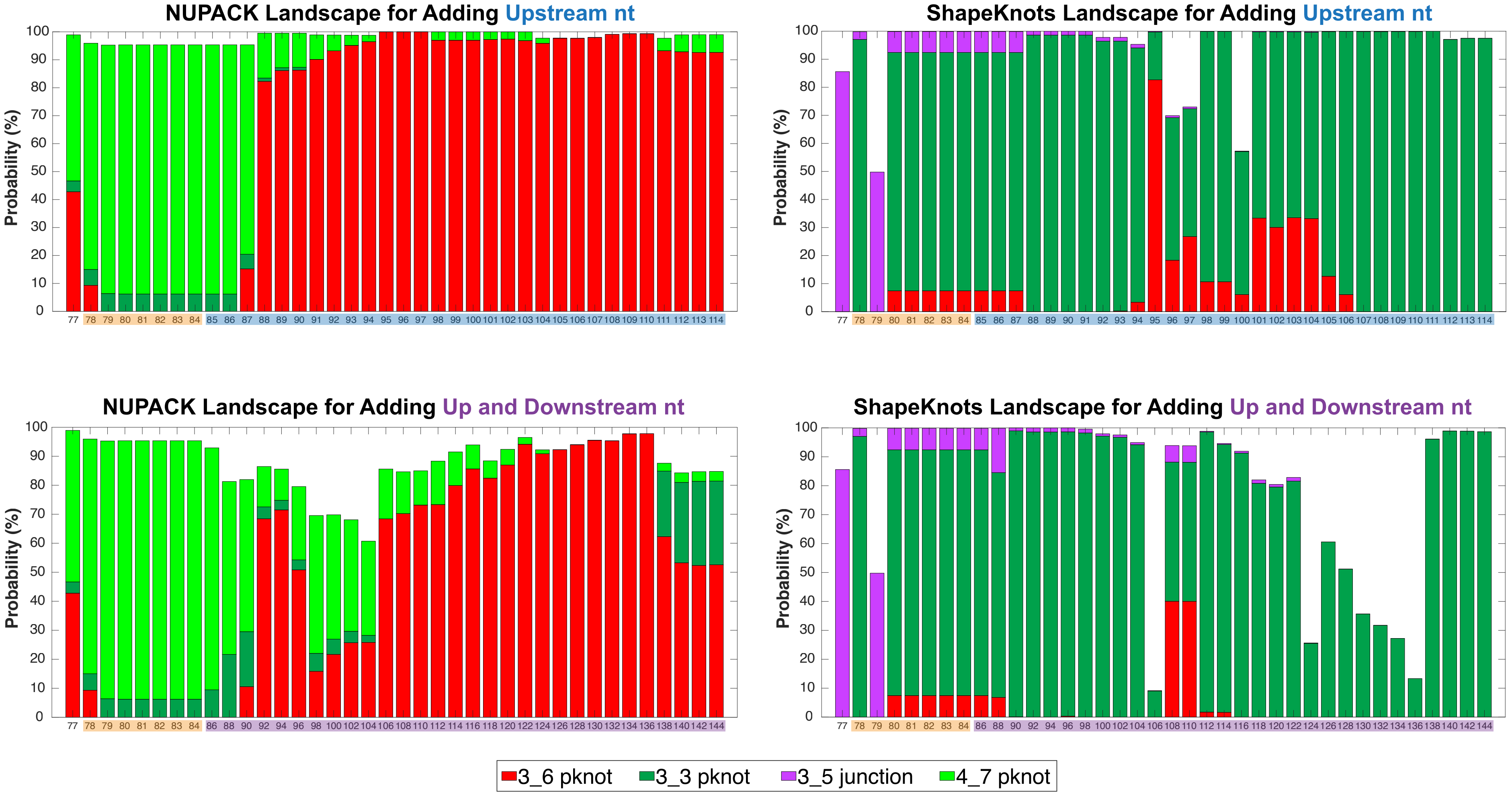


Figure S12: Conformational landscape of the frameshifting element for different sequence lengths predicted by NUPACK and ShapeKnots without any SHAPE reactivities. For each length, probabilities of all structures containing independently folded 3_6, 3_3, 3_5, and 4_7 are individually summed. The compositions are colored red (3_6), green (3_3), purple (3_5), and light green (4_7) respectively.

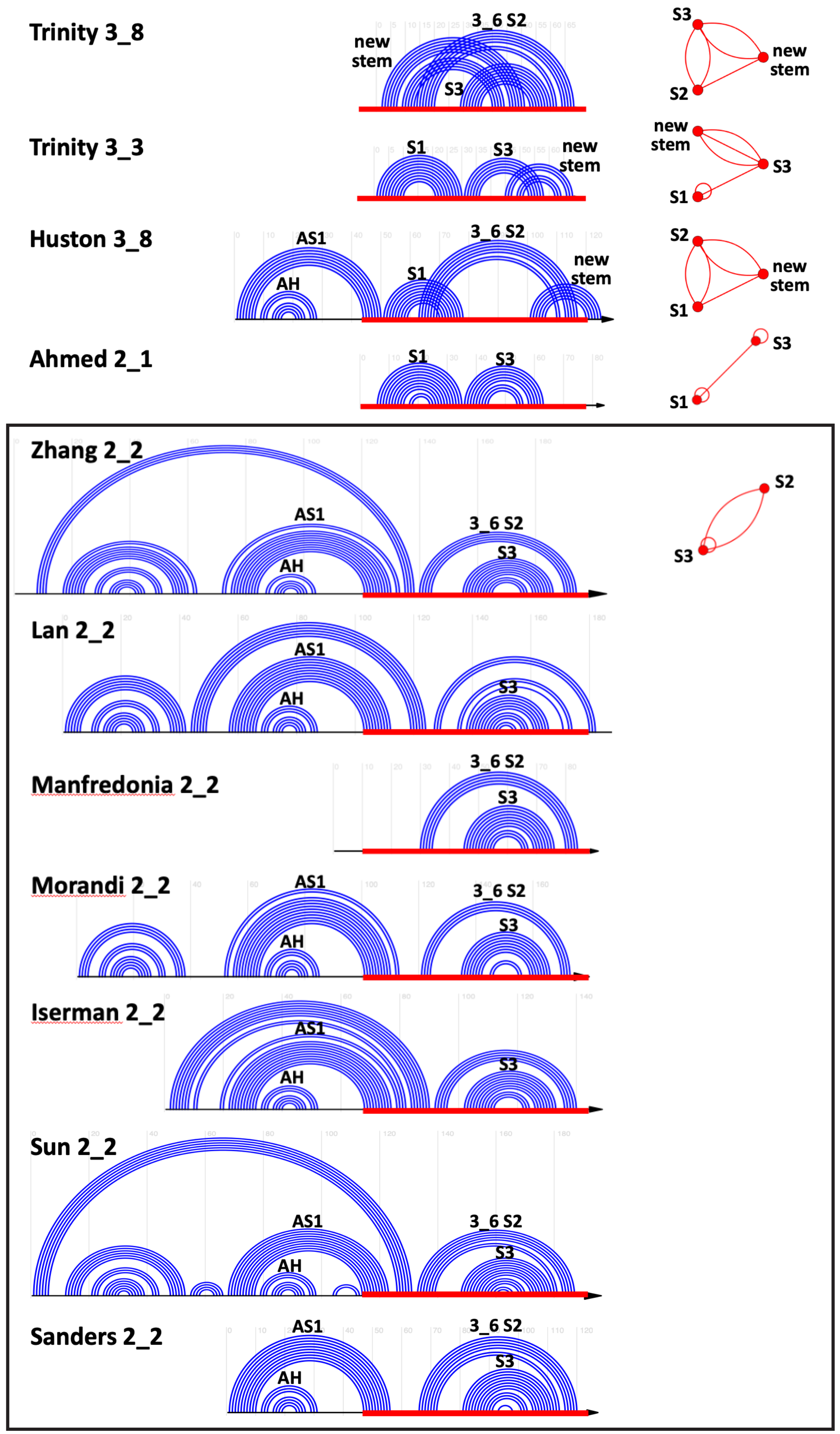


Figure S13: Other major and minor SARS-CoV-2 FSE conformations reported in the literature.[5–14] The common 77 nt FSE region is highlighted in red, and the sequences are aligned.

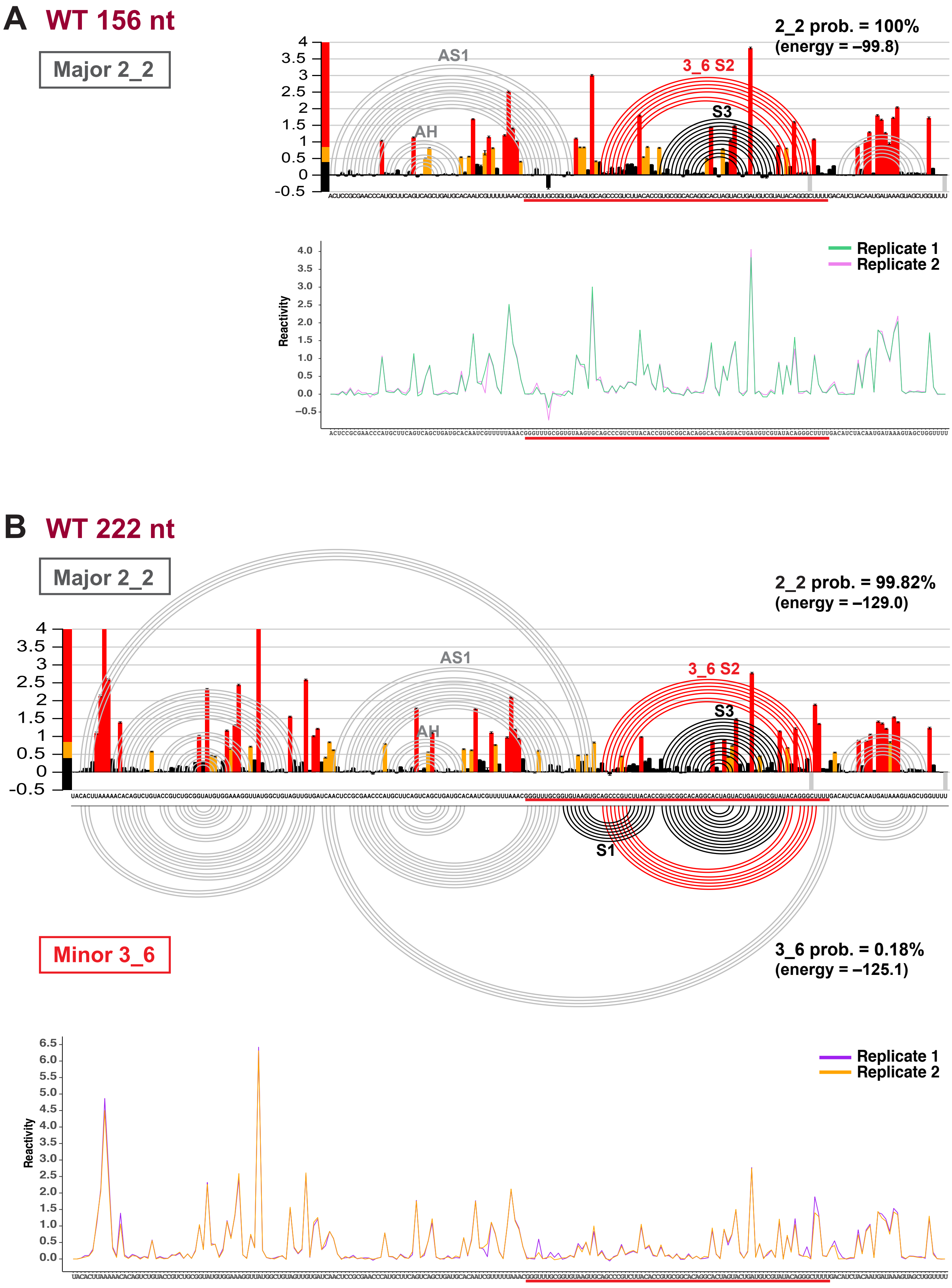


Figure S14: SHAPE reactivity analysis for wildtype (A) 156 nt and (B) 222 nt FSE constructs. For each construct, SHAPE reactivity and ShapeKnots predictions for Replicate 1 are shown at top, and alignment of two replicates at bottom. The common central 77 nt FSE region is highlighted in red.

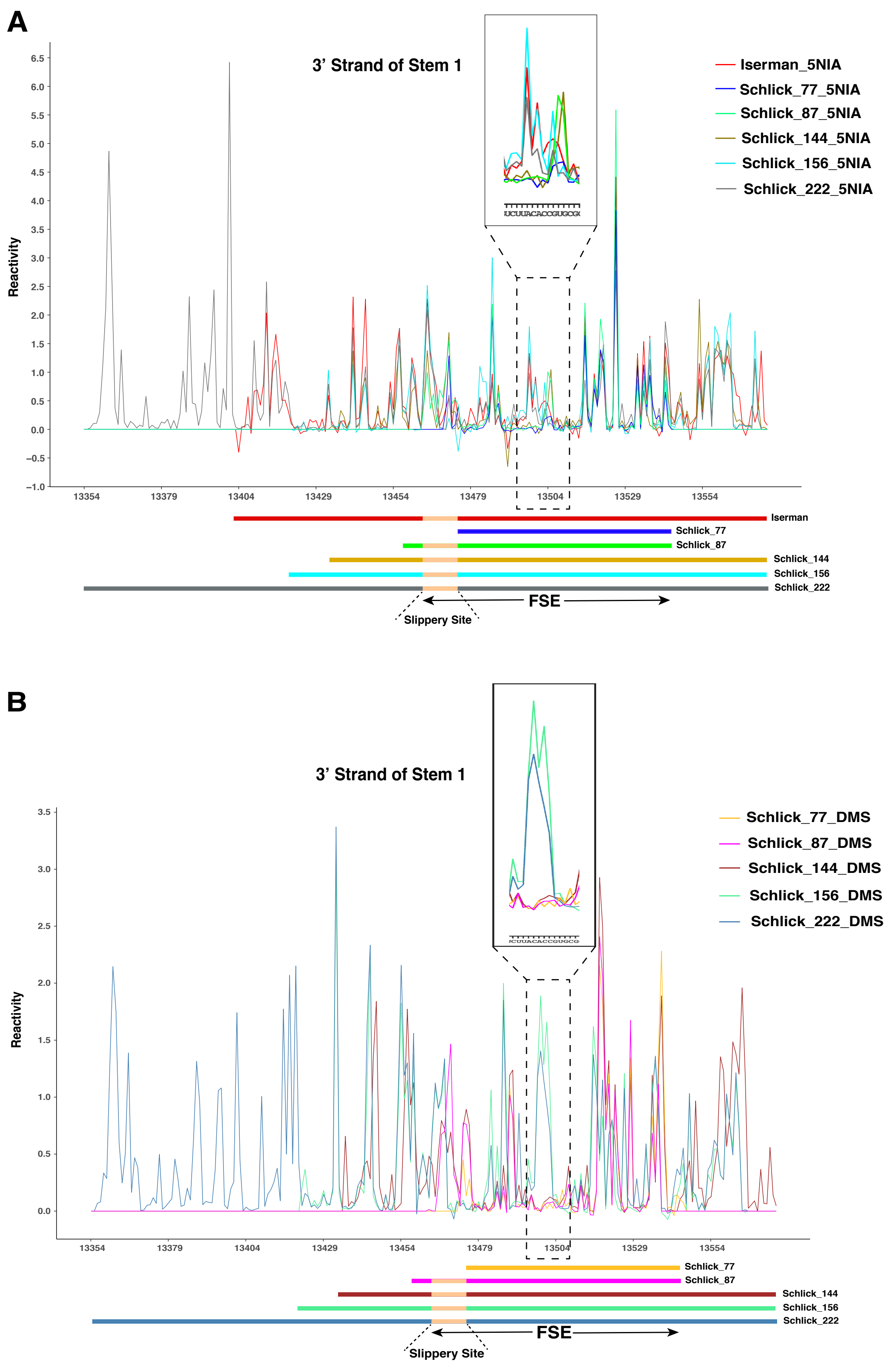


Figure S15: Comparisons of (A) SHAPE and (B) DMS probing between different FSE length segments (77, 87, 144, 156, and 222 nt) collected in this study. (A) Our SHAPE reactivity profiles of different lengths are aligned, and the Iserman et al.[12] profile is also included for comparison. All the constructs were *in vitro* transcribed and probed using 5NIA, and their schematic is shown at bottom. When the sequence length goes beyond 144 nt, a sudden increase in SHAPE reactivity is observed for residues 13495–13500, where the 3′ strand of Stem 1 locates, suggesting a key switch from Stem 1 to AS1. (B) DMS reactivity profile alignment, with a similar reactivity increase for residues 13495–13500.

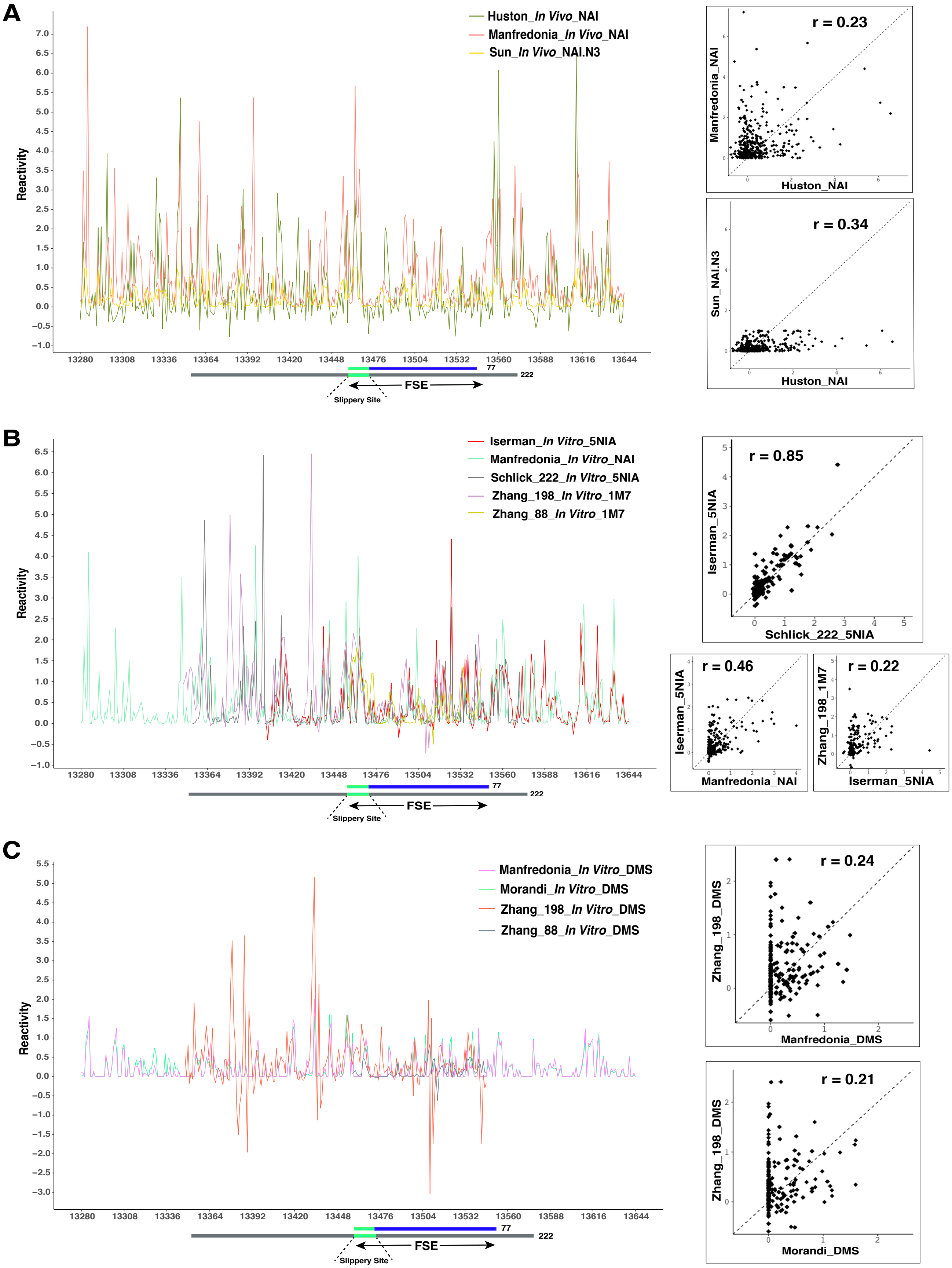


Figure S16: Comparisons of chemical probing by different groups for extended FSE region (residues 13280–13644). (A) Comparative *in vivo* genome-wide SHAPE reactivity profile from three groups.[6,10,13] Scatter plots on the right show low Pearson correlation coefficient ($r < 0.5$) between the datasets. (B) Comparative *in vitro* SHAPE reactivity profile from four groups including our 222 nt construct.[8,10,12] Low correlation between different datasets except for our 222 nt construct with Iserman et al. 1000 nt construct[12] ($r = 0.85$). (C) Comparative *in vitro* DMS reactivity profile from three groups,[8,10,11] again with low correlation.

### References


(1) Elbe, S.; Buckland-Merrett, G. Data, disease and diplomacy: GISAID's innovative contribution to global health. *Glob. Chall.* **2017**, *1*, 33–46.

(2) Katoh, K.; Misawa, K.; Kuma, K.; Miyata, T. MAFFT: a novel method for rapid multiple sequence alignment based on fast Fourier transform. *Nucleic Acids Res.* **2002**, *30*, 3059–3066.

(3) Antczak, M.; Zok, T.; Popenda, M.; Lukasiak, P.; Adamiak, R.; Blazewicz, J.; Szachniuk, M. RNApdbee – a webserver to derive secondary structures from pdb files of knotted and unknotted RNAs. *Nucleic Acids Res.* **2014**, *42*, W368–372.

(4) Petingi, L.; Schlick, T. Partitioning and Classification of RNA Secondary Structures into Pseudonotted and Pseudoknot-free Regions Using a Graph-Theoretical Approach. *IAENG Int. J. Comput. Sci.* **2017**, *44*, 241–246.

(5) Trinity, L.; Lansing, L.; Jabbari, H.; Stege, U. SARS-CoV-2 ribosomal frameshifting pseudoknot: Improved secondary structure prediction and detection of inter-viral structural similarity. **2020**, bioRxiv, doi: 10.1101/2020.09.15.298604, preprint posted September 2020.

(6) Huston, N. C.; Wan, H.; Strine, M. S.; de Cesaris Araujo Tavares, R.; Wilen, C. B.; Pyle, A. M. Comprehensive in vivo secondary structure of the SARS-CoV-2 genome reveals novel regulatory motifs and mechanisms. *Mol. Cell* **2021**, *81*, 584–598.e5.

(7) Ahmed, F.; Sharma, M.; Al-Ghamdi, A. A.; Al-Yami, S. M.; Al-Salami, A. M., et al. A Comprehensive Analysis of cis-Acting RNA Elements in the SARS-CoV-2 Genome by a Bioinformatics Approach. *Front. Genet.* **2020**, *11*, 1385.

(8) Zhang, K.; Zheludev, I. N.; Hagey, R. J.; Wu, M. T.-P.; Haslecker, R., et al. Cryo-electron Microscopy and Exploratory Antisense Targeting of the 28-kDa Frameshift Stimulation Element from the SARS-CoV-2 RNA Genome. **2020**, bioRxiv, doi: 10.1101/2020.07.18.209270, preprint posted July 2020.

(9) Lan, T. C. T.; Allan, M. F.; Malsick, L. E.; Khandwala, S.; Nyeo, S. S. Y., et al. Insights into the secondary structural ensembles of the full SARS-CoV-2 RNA genome in infected cells. **2021**, bioRxiv, doi: 10.1101/2020.06.29.178343, preprint posted February 2021.

(10) Manfredonia, I.; Nithin, C.; Ponce-Salvatierra, A.; Ghosh, P.; Wirecki, T. K., et al. Genome-wide mapping of SARS-CoV-2 RNA structures identifies therapeutically-relevant elements. *Nucleic Acids Res.* **2020**, *48*, 12436–12452.

(11) Morandi, E.; Manfredonia, I.; Simon, L. M.; Anselmi, F., et al. Genome-scale deconvolution of RNA structure ensembles. *Nat. Methods* **2021**, *18*, 249–252.

(12) Iserman, C.; Roden, C. A.; Boerneke, M. A.; Sealfon, R. S.; McLaughlin, G. A.; Jungreis, I.; Fritch, E. J., et al. Genomic RNA Elements Drive Phase Separation of the SARS-CoV-2 Nucleocapsid. *Mol. Cell* **2020**, *80*, 1078–1091.

(13) Sun, L.; Li, P.; Ju, X.; Rao, J.; Huang, W.; Ren, L.; Zhang, S., et al. In vivo structural characterization of the SARS-CoV-2 RNA genome identifies host proteins vulnerable to repurposed drugs. *Cell* **2021**, *184*, 1865–1883.e20.

(14) Sanders, W.; Fritch, E. J.; Madden, E. A.; Graham, R. L.; Vincent, H. A.; Heise, M. T.; Baric, R. S.; Moorman, N. J. Comparative analysis of coronavirus genomic RNA structure reveals conservation in SARS-like coronaviruses. **2020**, bioRxiv, doi: 10.1101/2020.06.15.153197, preprint posted June 2020.
